# Integration of polygenic risk with single cell methylation profiles for depression

**DOI:** 10.1101/2025.03.11.642615

**Authors:** Xinzhe Li, Kangcheng Hou, Katherine W. Eyring, Cuining Liu, Chongyuan Luo, Daniel H. Geschwind, Bogdan Pasaniuc

## Abstract

Large scale genome-wide association studies (GWAS) have identified hundreds of risk loci for major depression disorder (MDD) with their functional understanding being largely unknown. We integrate MDD polygenic risk from GWAS with methylation at a single cell level resolution to gain insights into the role of methylation in driving MDD risk. We introduce a new approach that leverages the polygenic risk of disease with single-cell methylation data to provide a methylation single cell disease relevance score (met-scDRS) for every cell in a single-cell methylation-seq experiment. We analyzed human atlas single cell methylation data to find 54.0% of layer 2/3 intratelencephalic (L2/3-IT) neurons and 46.5% of layer 5 extratelencelphalic (L5-ET) neurons in the dataset showing significant met-scDRS enrichment. We identified gradient of met-scDRS from inferior temporal gyrus to middle temporal gyrus and variations in posterior to anterior brain axis within L2/3-IT neurons. Met-scDRS identifies functional pathways such as synaptic cellular component, somato-dendritic compartment, post-synapse, cell junction organization that are implicated in diseases and identifies genes that are more disease associated. We contrasted met-scDRS for MDD across 75 other traits including brain, immune/blood, metabolism, and other trait categories to identify diverging and converging cell types and prioritized pathways across different traits. Finally, we demonstrated that met-scDRS is portable across non-CpG and CpG methylation data in providing robust signal.

## Introduction

Cytosine methylation is a covalent modification of DNA that is found in most CpG dinucleotides in the mammalian genome^1^. In most somatic mammalian tissues, over 70% of CpG sites are methylated while unmethylated CpG sites are enriched in promoters and distal regulatory elements^2^. Exceptionally high levels of non-CpG methylation, or methylation of cytosines outside of CpG dinucleotides, accumulate in human brain tissues during post-natal development^3^. The genome-wide pattern of non-CpG methylation is correlated with gene expression^2,4^ and is known to have functional importance in gene silencing^2,5,6^, gene environment interaction^7^, cell type specificity^8,9^, and disease association^2,10,11^. There are few methodologies that integrate GWAS with methylation single cell data at single cell resolution, with most methods focusing on finding heritability in differentially methylated regions^10^ or comparing cell-cell methylation differences between cases and controls^12^. Not only do they not leverage GWAS, they often disregard non-CpG methylation, which may have different characteristics from CpG methylation^2,4,8^. For identifying context, such as cell types relevant to disease, current methods focus on utilizing pre-existing marker genes and quantifying their associations with existing GWAS^13,14^ or directly comparing proportional differences between case and control^15–18^. Although tools such as single-cell disease relevance score^19^ (scDRS) and sc-linker^20^ exist to integrate GWAS and transcriptomics at single cell level resolution, they are built with RNA sequencing modality in mind and do not extend to other genomic modalities. Combined, these methods do not provide single cell resolution and may not be sufficient in understanding the whole landscape of signal heterogeneity in single cell methylomes.

Recent technological developments generate single cell methylome datasets^21–24^ that resolve cell types, probe disease associated genomic regions, and methylation related gene expression changes. Such datasets provide a valuable opportunity to integrate single cell methylome with GWAS and allow for better mechanistic view on how methylation modification at putative disease genes could confer disease risks in a context dependent manner. We introduce met-scDRS, an approach that integrates non-CpG and CpG methylation with GWAS at single-cell level resolution to provide a methylation single cell disease relevance score(met-scDRS) for every cell in the assay. Using GWAS genes and cell level control gene sets that have matched variance and methylation levels from the single cell methylome, met-scDRS tests for hypomethylation of disease risk genes compared to genome wide averages sampled from those control gene sets.

We used simulation to find that met-scDRS is well calibrated when no disease association is present in the data while also being well powered when a fraction of cells in the experiment are disease implicated. We used met-scDRS on a MDD GWAS^25^ with the non-CpG human methylation atlas data^22^ to compute cell-specific disease scores for finding disease associations with different cell-types and spatial regions of the brain. In line with previous works that shows MDD association with L2/3-IT glutamatergic neurons^26–28^, we found that over 50% of L2/3-IT glutamatergic neurons are significantly associated with MDD(< 0.1 FDR corrected). We identified a significant cell type specific dissection region effect, demonstrating that dissection region significantly predicts risk score within cell type context. Within the L2/3-IT neuronal type, we observed a gradient of met-scDRS in temporal gyri as well as substantial variations along the posterior to anterior brain axis. When applied to CpG methylation, met-scDRS was able to identify over 30% of L2/3-IT excitatory neurons which are significantly associated with MDD while uncovering CpG methylation specific cell types in the disease. Next, we used met-scDRS as a gene / pathway prioritization tool to discover functional pathways in non-CpG methylation for MDD. We found gene sets with specific and functional roles with neurons, such as synaptic cellular components and neurotransmitter pathways where both of these pathways have well established important role in MDD^29–33^. Finally, we contrasted MDD scores with met-scDRS across 75 traits spanning brain, blood/ immune, metabolism and other related traits in both sequencing modalities and found that met-scDRS identified disease relevant cell type – trait pairs.

Overall, we showed that met-scDRS is a useful tool for integrating GWAS with single-cell methylation data to identify disease association heterogeneity in cell type and to gain insights into biological basis of depression and other polygenic traits.

## Results

### Overview of method

Met-scDRS takes as input a single-cell methylation dataset and a disease gene set (from GWAS) to compute a disease score/p-value for each cell in the data (**Figure 1**). To account for expected anticorrelation between gene expression and raw methylation level^23^, met-scDRS computes the inverse fraction X*^’^_c,g_* of the raw methylation level *X_c,g_* as 1-X where the raw methylation level is defined as methylcytosine base coverage divided by the total cytosine base coverage aggregated at gene bodies (**Methods**). For CpG methylation, to account for the fewer number of CpG sites in the genome^34^, met-scDRS aggregates X*^’^_c,g_* for each cell with 5 neighboring cells in UMAP space. First, control genes with similar methylation level and variance as the disease putative genes from X*^’^* are identified. Second, met-scDRS computes a polygenic disease score for each cell for all gene sets by computing the variance weighted product between gene weights and methylation level. Third, met-scDRS computes an empirical p-value by comparing the disease score to the control set (**Methods**)^19^.

**Figure 1:**
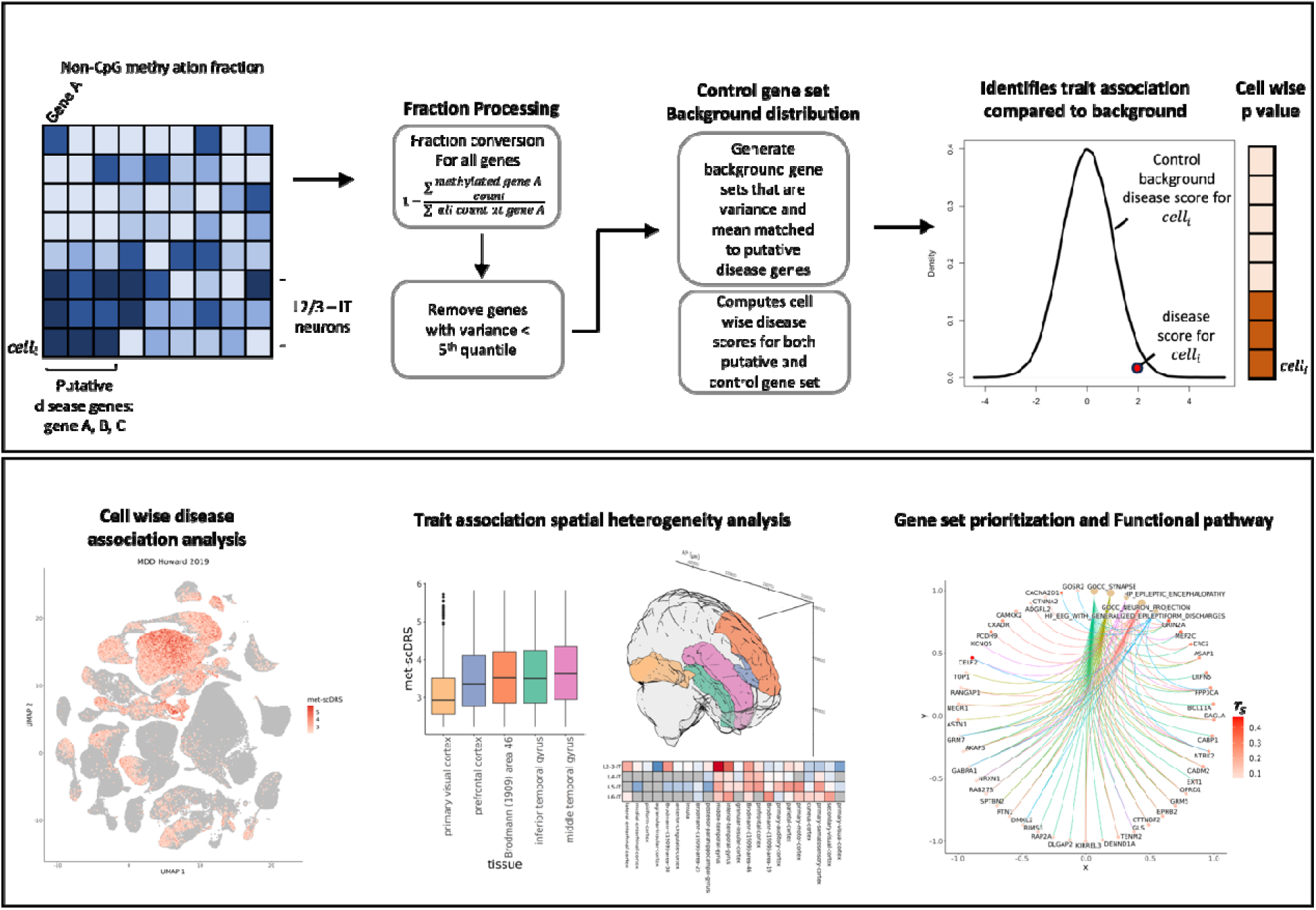
(top) Met-scDRS algorithm, from left to right: gene level methylation fraction first reversed by 1 – original fraction level for consistency in directionality with respect to gene expression. Genes with very low variance are filtered for variance stabilization. Using the putative gene set, met-scDRS generate control gene set with matching methylation fraction and variance level and computes cell-wise disease risk score and p values. (bottom) Example of downstream analysis overview from this work. From left to right: UMAP visualization on met-scDRS risk score; cell type specific spatial heterogeneity analysis across different brain regions; gene ontology pathway analysis by using prioritized gene set from met-scDRS.

### Met-scDRS is calibrated in simulations

We investigated the calibration of met-scDRS under different simulation scenarios: random gene selection, highly hypomethylated gene selection, and high variance gene selection (**Methods**). These three scenarios are selected to assess met-scDRS robustness at 1) random genes 2) hypomethylated genes and 3) genes with high methylation variance. Met-scDRS is well calibrated in all scenarios and remains unbiased with different gene set sizes for all simulated scenarios (**Figure 2a, Supplementary Figure 1, 2, 3, 4**). Next, we investigated how statistical power changes with varying proportions of false positive disease genes within the input gene set as well as with increasing hypomethylation effects in disease genes (**Methods**). We found that met-scDRS power increases linearly with increasing true positive genes in the disease gene set (**Figure 2b**). Met-scDRS is well powered (> 75%) even when the perturbed effect size is low (< 0.01 times fraction) as power grows exponentially with perturbations (**Figure 2c**). In line with our calibration simulation studies, we found that the variance between causal simulations is also low for all scenario and replicates (**Figure 2, Supplementary Figure 1, 2, 3, 4)**. Combined, our simulation studies show that our method is well powered to identify cells that are causally associated with disease while being well calibrated when there are no signals associated with the data.

**Figure 2:**
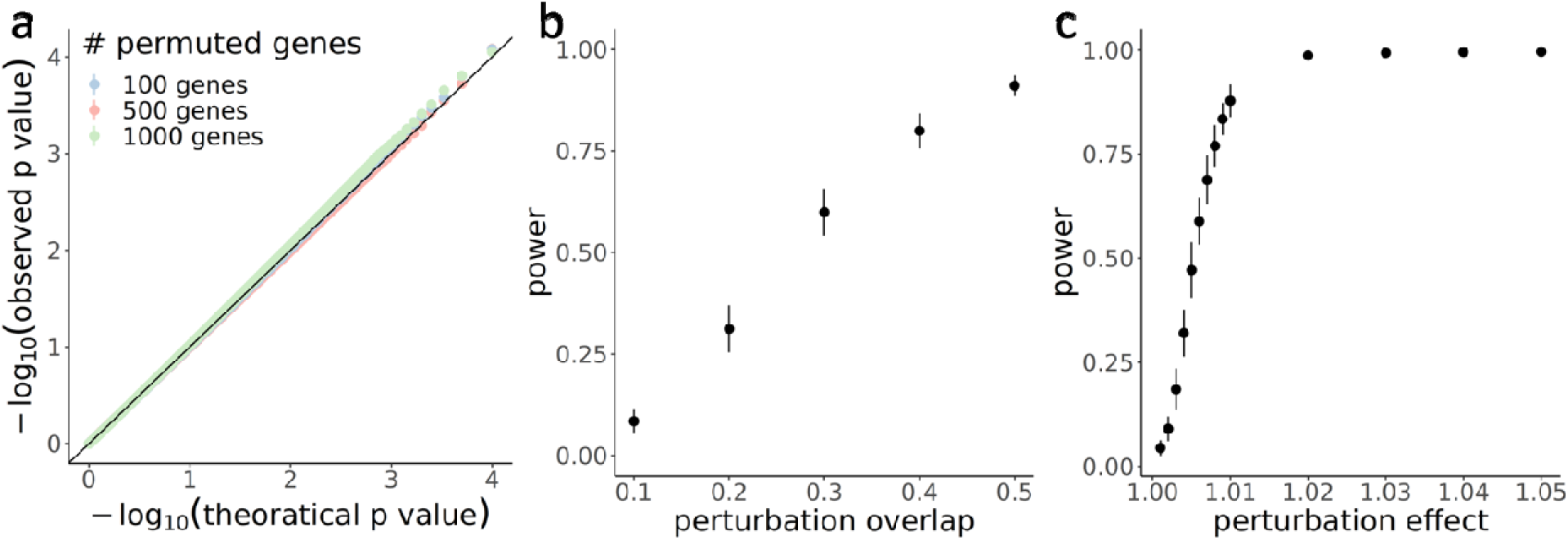
Met-scDRS is calibrated in simulations. (a) Calibration curve showing observed p values against the theoretical p values colored by different number of putative disease genes randomly selected from a set of genes with > 95th percentile in variance. (b) Power curves showing power computed with overlap from 10% - 50% of the GWAS putative disease genes selected as causal genes. (c) Perturbation effect from 1.001 – 1.05 times of the original fraction. All confidence intervals are computed as 2 * standard deviation using 100 replicates.

### Met-scDRS identifies disease relevant cells in human methylation atlas for MDD

Next, we applied met-scDRS on a publicly available human methylation atlas data^22^ sequenced across 46 brain regions and 188 quantified cell types generated from adult human brain using MDD genes derived from GWAS^25^. We identified a total of 50,332 cells with significant met-scDRS (< 0.1 FDR corrected) out of 373,888 cells in the data. 92.5% of the significant cells are glutamatergic excitatory neurons (n = 46,537) followed by a small number of inhibitory neurons and non-neuronal cells. Within the significant excitatory neurons, L2/3-IT neurons are the most abundant (n = 30,623), followed by L5-IT neurons (n = 4,588) and L6-IT neurons (n = 3,014) (**Supplementary Table 1**). Together, they represent 54.0%, 26.2% and 26.7% of all the cells within their respective cell type (**Supplementary Table 2**). This suggests that both superficial layer projecting neurons and deep layer projecting neurons are implicated in MDD to different extents. For inhibitory neurons, Lamp5 and medium spiny neurons expression D2 receptor (MSN-D2) are amongst the most implicated inhibitory neurons (n = 1,037 and 956 respectively, representing 9.71% and 5.71% of cells in cell type). Compared to the other cell class, non-neuronal cells make up the least number of cells significantly associated with MDD (n = 147 out of 34,722 total non-neuronal cells) (**Figure 3a**), likely due to the low non-CpG methylation presence in non-neuronal cells.

**Figure 3:**
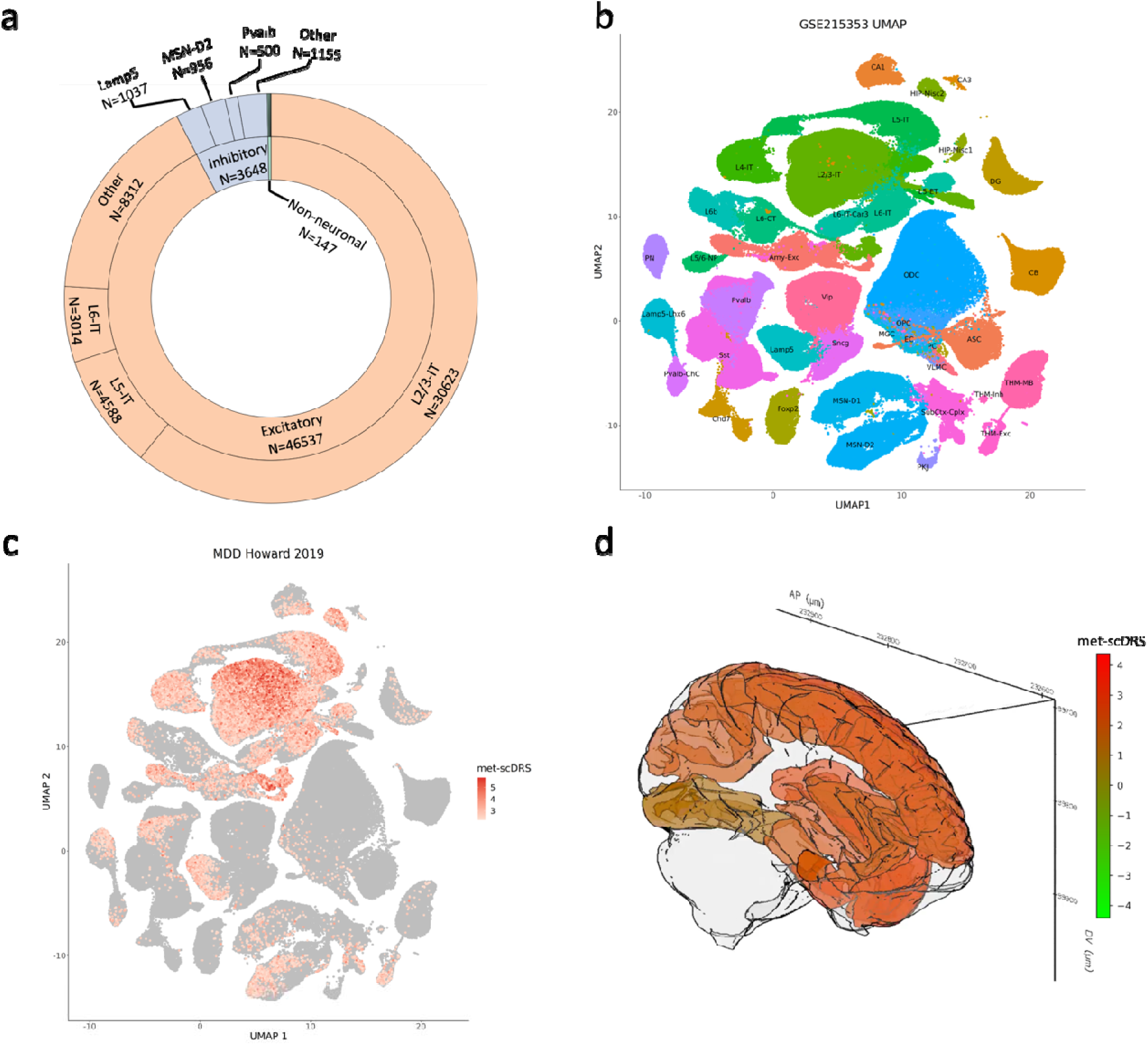
Analysis overview for human methylation fraction dataset using MDD GWAS genes as putative disease genes. (a) Pie chart showing the break-down of cell class, cell type identities for cells with significant association with MDD. Pie chart is colored by cell class and labeled with number of significant (FDR corrected p value < 0.1) cells that belong to each of the categories. Only the top 3 cell types for excitatory neurons and inhibitory neurons are shown for simplicity. (b-c) UMAP of the human methylation atlas colored by (b) cell type identities and (c) met-scDRS risk score computed with non-CpG modality in MDD where the significant cells are colored with a gradient from white to red by magnitude of met-scDRS and the non-significant cells are colored in grey. (d) spatial render of brain regions colored by average MDD met-scDRS on non-CpG modality at each of region in common cortex axis for L2/3-IT neurons. Color scale defined by range of MDD met-scDRS from minimum (green) to maximum (red)

To validate that met-scDRS is not simply prioritizing cell types that have higher global methylation levels, we visualized the non-CpG methylation distribution in each of the cell type across excitatory neurons, inhibitory neurons and non-neuronal cells (**Supplementary Figure 5-7**). We observed non-significant correlation between average non-CpG methylation and proportion of significant cells across cell type (spearman correlation = −0.19, p value = 0.23). To validate that our findings are not driven by batch effects, we visualized the number of significant cells by batch and donors and identified no donor effect but some relationship with sequencing batch (**Supplementary Figure 8, 9**). For stringency, we regressed out batch effects (**Methods**) and observed highly correlated results pre and post batch effect correction (spearman correlation > 0.99) (**Supplementary Figure 10**). Collectively, met-scDRS prioritizes glutamatergic neurons, particularly the L2/3-IT neuronal type as the main associated cell type toward MDD.

Next, we investigated variation in met-scDRS within cell types and found a gradient of scores within L2/3-IT and L5-IT excitatory neurons. This suggests that a subset of cells within these two neuronal types are particularly hypomethylated in MDD putative genes and are more disease implicated. We first investigated if there are subtypes within L2/3-IT and L5-IT neurons that could explain such variation. Indeed, in both neuronal types, the number of significantly associated cells is not uniformly distributed across subtypes. In L2/3-IT neurons, subtypes 1-5 and 7-11 have higher proportion in significant cells compared to the rest of the subtypes (**Supplementary Figure 11a, 11b, 12**) whereas in L5-IT neurons, subtype 5 is enriched in significantly associated neurons (**Supplementary Figure 13a, 13b, 14**). Since the subtypes are significantly (*simulated X*^2^ *p value < 5 x 10^-^*^6^) associated with dissection tissue region in both of the L2/3-IT and L5-IT cell types, we reason that within-cell type regional differences could explain the granularity in risk. To this end, we visualized the average region-wise disease risk score within L2/3-IT neurons on a 3D human brain model in the common cortex axis (**Figure 3d**, **Supplementary Table 3**). Overall, most of the brain’s physical dissection regions show moderately high average associations (met-scDRS > 2), with a large deviation between regions that ranges from a minimum at hippocampus field (average met-scDRS of 0.778) to a maximum at lateral-entorhinal region (average met-scDRS of 3.08). After quantifying the specificity index for each region and cell type pair, we found that the origin of intratelencephalic neurons along common regional axis of the cortex show varying levels of MDD risk predictability in a cell type specific manner (**Methods**, **Figure 4a**). Specifically, origin of intratelencephalic projecting neurons from prefrontal cortex, Brodmann area 46, inferior temporal gyrus, and middle temporal gyrus are predictive of risk association with MDD to varying degree (specificity index mean = 1.07) with regions, such as Brodmann area 19, being more specific to L6-IT neurons in risk association (specificity index =1.47) (**Supplementary Table 4**). A stronger association with MDD is present for L2/3-IT neurons derived from middle temporal gyrus than for those in the inferior temporal gyrus (**Figure 4a, c**). This demonstrates a ventral to dorsal gradient in risk association to MDD for L2/3-IT neurons among temporal gyri. Investigating how the risk score varies in a cell type specific manner across anterior to posterior axis, we found that the met-scDRS by regions for L2/3-IT neurons broadly increase across the primary visual cortex, Brodmann region 44-45, Broadman region 46, inferior temporal gyrus and middle temporal gyrus (**Figure 4b**).

**Figure 4:**
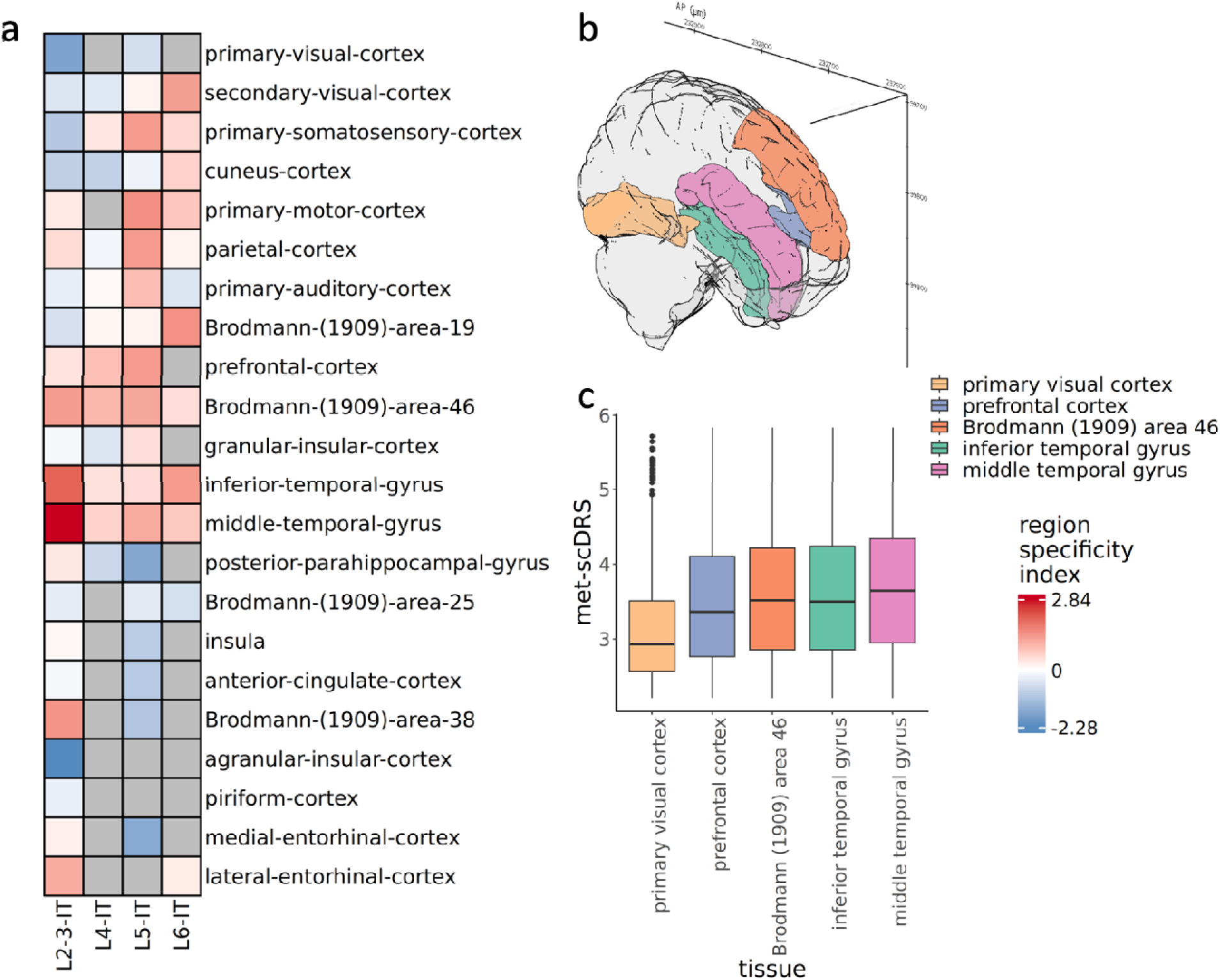
(a) Heatmap colored by region specificity index computed in non-CpG in MDD for layer L2/3, L4, L5, and L6 intratelencephalic projecting neurons across common regional axis. (b) 3D rendering of brain models colored by primary visual cortex (yellow), prefrontal cortex (blue), Brodmann region 46 (orange), middle frontal cortex (purple), inferior temporal gyrus (green). (c) Boxplot on the non-CpG met-scDRS for different regions with same coloring scheme as brain rendering.

### Met-scDRS prioritizes functional genes

Given the strong signal in MDD association at a cell-type resolution, we hypothesized that groups of genes in the putative gene set could belong in the same biological pathway and collectively confer risk via methylation. The top 100 genes with the most contribution to the MDD met-scDRS are strongly enriched in synaptic related pathways (**Figure 5a**). Synaptic cellular component, somatodendritic compartment, post-synapse, and cell junction organization are among the most over-represented pathways (FDR adjusted p < 0.01) (**Figure 5a**). We also visualized the membership of genes in significant pathways colored by the spearman correlation for each gene member. Among these genes, CELF2, a gene well known for its neuronal differentiation regulatory role^35^, had the highest spearman correlation coefficients (rho = 0.47) (**Figure 5b**) between their methylation and risk score. GRIN2A and CACNA2D1, two protein coding genes for ion channels that are critical to establish synaptic transmission^36,37^, are also moderately correlated between its methylation level and met-scDRS (**Figure 5b**) with spearman correlation of 0.36 and 0.28 respectively. Together, met-scDRS identifies genes with hypomethylation status involved in synaptic establishment and transmission is particularly implicated in MDD.

**Figure 5:**
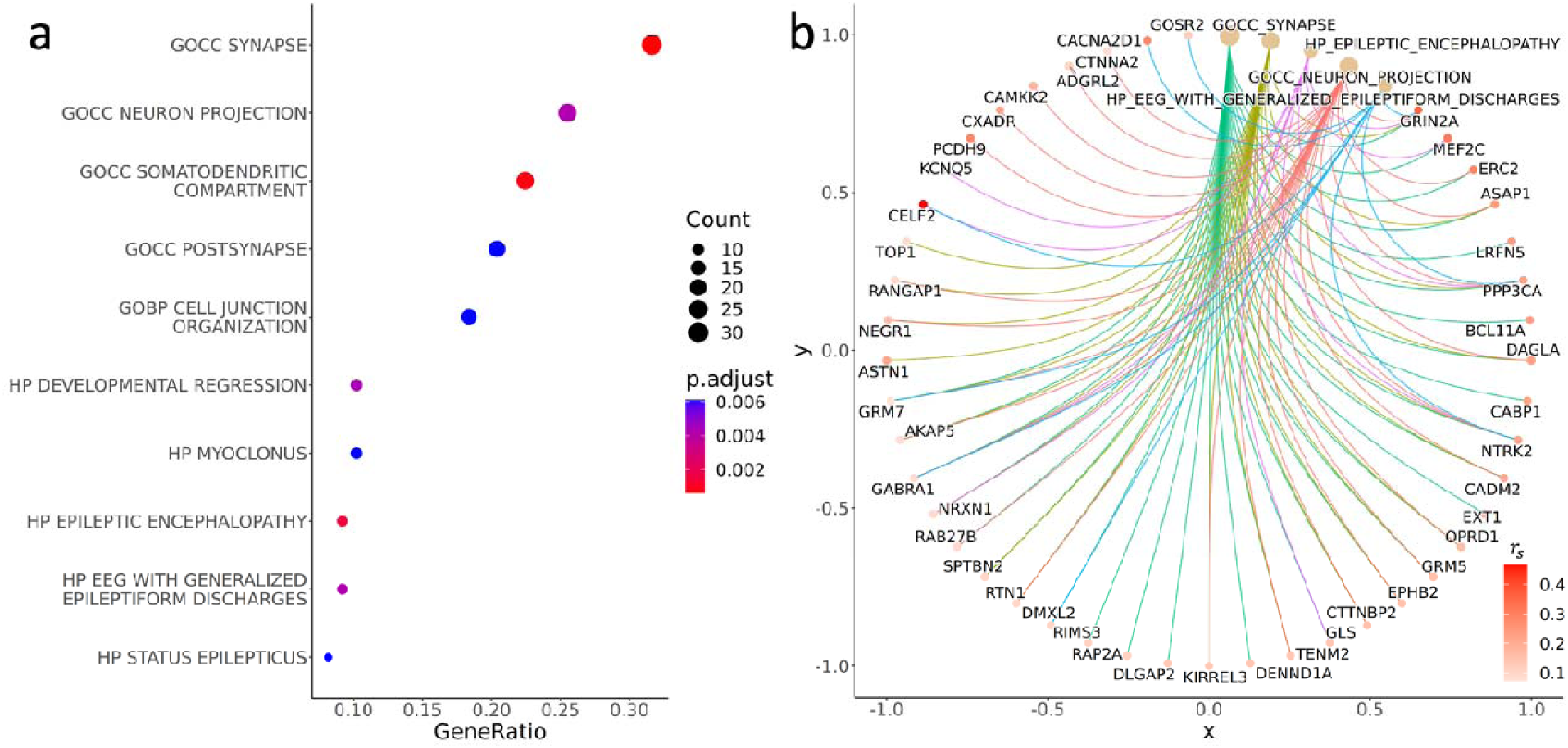
(a) Gene ontology analysis using met-scDRS prioritized gene set as foreground set and MDD GWAS genes as background set. For each of the pathways on the Y axis, X axis shows the gene ratio between foreground and background; each dot is colored by FDR adjusted p value and sized by the number of genes in foreground that belong in that pathway. (b) Network diagram that shows membership of genes that belong in each of the pathway enriched for MDD with lines colored by the pathways and dots colored by spearman correlation between the processed methylation fraction and met-scDRS risk score.

### Met-scDRS detects divergent and convergent cell types associated with brain traits

Finally, we expanded analyses to 75 traits broadly spanning five categories: brain traits, immune / blood traits, heart traits, metabolic traits, and other traits (**Supplementary Table 1, 2, Supplementary Figure 15**). Since the single-cell atlas consists of only brain samples, we hypothesized that met-scDRS will only show a high number of disease-associated cells in the brain related traits. Indeed, the average percentage of significant cells (FDR < 0.1) associated with diseases in each of the five disease categories was enriched for brain-related traits: 26,443 (7.07%), 5,000 (1.34%), 3,824 (1.02%), 1,487 (0.40%) and 730 (0.20%) significant cells are associated with brain, metabolic, other, blood/immune, and heart related traits respectively. Next, we quantified for the three cell classes (excitatory neurons, inhibitory / subcortical neurons, and non-neuronal cells), the average percentage of significant cells in each of the three classes across the five types of traits. Brain traits have the highest proportion and variance of significant cells in excitatory neurons, suggesting an abundance of disease associations between hypomethylation and brain related traits in excitatory neurons with substantial variation within brain traits (**Figure 6a**). In contrast, blood/immune traits, heart traits, metabolic traits and other traits all have minimum proportion of significantly associated excitatory neurons (mean percentage < 5%), showing that met-scDRS identifies excitatory neurons as disease relevant cell types associated to brain traits given the dataset. We interpret the result as met-scDRS detects hypomethylation in excitatory neurons of brain samples as the primary source of signal in brain traits and are well calibrated to not detect significant association in non-brain traits while suggesting substantial variation within brain traits and within excitatory neurons (**Figure 6 a, b**). Cell types such as L2/3-IT, L5-ET, L6-IT, and HIP-Misc2 neurons that are broadly associated across many traits, with substantial variation in proportion across traits (**Figure 6b, Supplementary Table 2**). Meanwhile, some cell types are more specific in association with diseases such as L6-IT-Car3, L6b, L6-CT neuronal types. We found SCZ to be overrepresented in the significant cells of excitatory deep layer projecting neurons such as L6-CT, L6b neuronal type as well as inhibitory neuronal type such as MSN-D1, and MSN-D2 compared to MDD. In contrast, MDD shows a higher proportion of significant cells in shallower regions projecting excitatory neuronal type such as L2/3-IT projecting neuron and hippocampus miscellaneous excitatory neurons.

**Figure 6:**
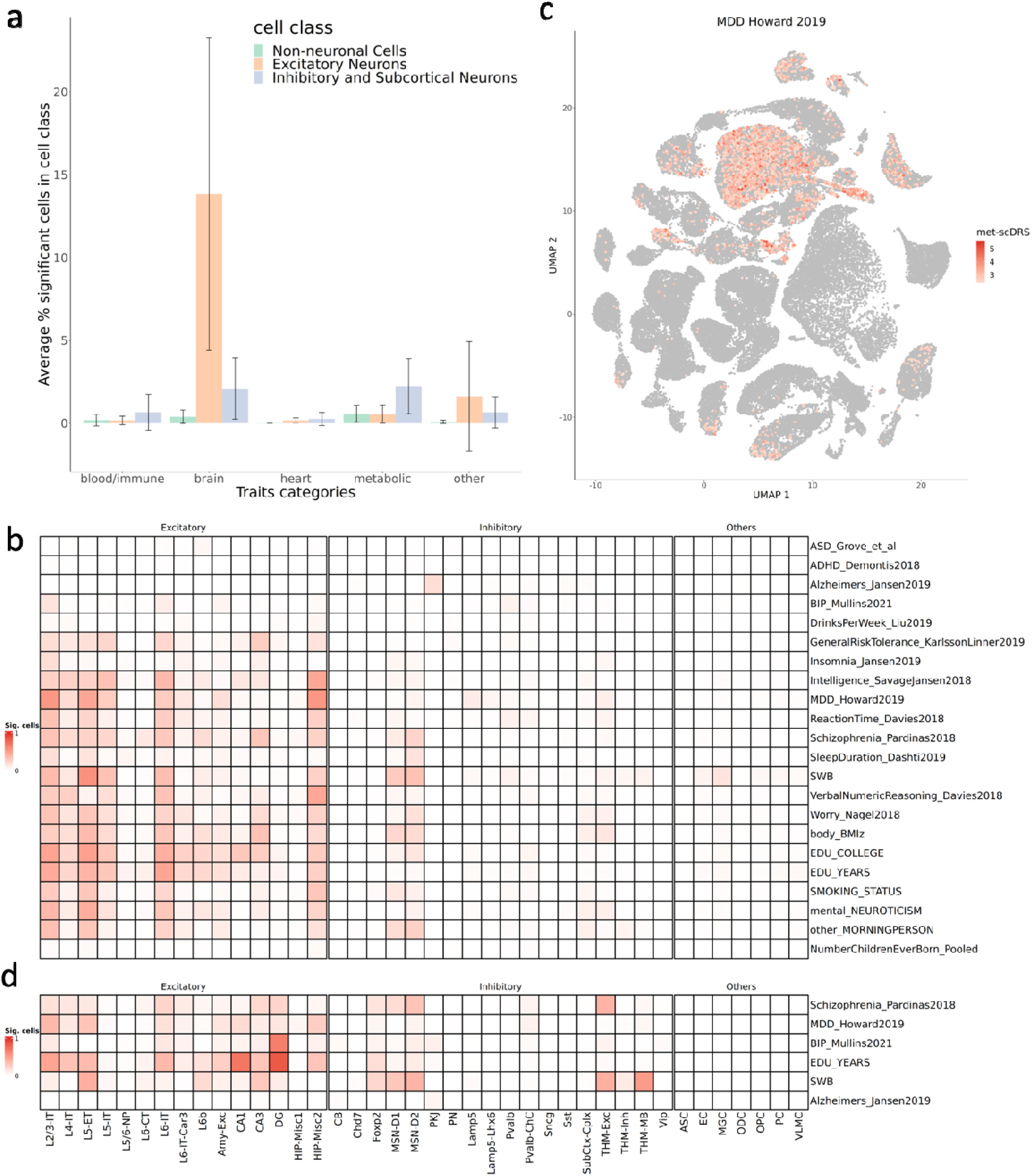
Bar chart (a) showing the average percentage of cells that are significant for each trait categories colored by each cell class. Confidence interval shows standard deviation. (b) Heatmap showing the proportion of significant cells in each of the cell type across brain related traits for non-CpG modality. Cell types are partitioned by excitatory (left), inhibitory neurons (middle) and non-neuronal cell types (right). (c) Met-scDRS risk score on UMAP computed using CpG modality where the significant cells are colored with a gradient from white to red and the non-significant cells are colored as grey. (d) Same as (b) but for CpG methylation modality for selected brain traits.

Next, we also investigated if the subtypes within the major cell types exhibit the same pattern where some subtypes maybe broadly associated across brain traits and some maybe more specific. Indeed, across a total of 16 L2/3-IT subtypes, we identify subtypes that are significantly associated in MDD such as subtype 1, 5, 11 which are also observed in SWB, neuroticism, and education traits (**Supplementary Figure 12**). However, some subtypes also show variation between diseases. For example, > 60% of neurons in subtype 8-10 are significantly associated with MDD, but only < 22% of neurons in subtype 8-10 are significantly associated with bipolar disorder. Interestingly, we see the same pattern in L5-IT neuronal subtypes, where 84.2%, 66.7% and 6.1% of neurons in subtype 5 is significantly associated with SWB, MDD and bipolar respectively (**Supplementary Figure 14**). Such patterns suggests that different cell types and subtypes show convergence and divergence across disorders and met-scDRS detects these homo-/heterogeneities in disease specificity. Additionally, we also found that some diseases such as Alzheimer’s have a lower proportion (< 5% significant cells) across most cell types and subtypes, revealing the possibility that hypomethylation may not be involved in pathogenesis or the dataset may not have captured relevant cell types. To ensure that the patterns we saw are not driven by non-CpG methylation expression, we again computed the correlation between non-CpG methylation and proportion of significant cells in the cell type. We found only one trait (number of children ever born) that has a significant spearman correlation after correction (spearman correlation = −0.56, FDR corrected p value = 0.014). For completeness, we also regressed out batch effect and observed met-scDRS having high correlation before and after correction (average spearman correlation > 0.98) across disorders.

### Met-scDRS gains insights into heterogeneity of cell-specific signals

We reasoned that there could be sets of common and specific biological pathways that maybe enriched across disorders, which would explain the convergence and divergence of trait association across cell types. We found common pathways that are enriched among brain traits such as the cellular component synaptic pathways (17 / 22 traits), synaptic signaling (11 / 22 traits), somatodendritic pathways (11 / 22 traits) (**Supplementary Table 5**). These common pathways suggest that there might be a shared disease etiology related to hypomethylation, which warrants further disease mechanistic studies. There are also pathways that are more disease specific. One example involves MDD, which shows strong enrichment in cellular component dendritic tree pathways, a pathway only significantly enriched in 6 out of 22 brain traits. Another example involves social well-being and insomnia where ion channel related pathways show strong enrichment compared to the rest of the brain related traits, suggesting insomnia and social well-being could share similar disease etiology (**Supplementary Figure 16a, b, Supplementary Table 5)**. Importantly, the social well-being prioritized gene set by met-scDRS is significantly (FDR p < 0.05) enriched in multiple ion channel related pathways, demonstrating strong pathway specificity (**Supplementary Figure 16a**). The pathways identified are functionally specific, which points to cellular phenotypic altering genes as these ion channels are fundamental in determining the action potential of neurons and their subsequent firing patterns^38^. This result hints that the variation in hypomethylation at ion transporters in neuronal cells is specifically important in the social well-being phenotype. For bipolar disorder, regulation of nervous system processes and somatodendritic compartment components are among the most significant (FDR p < 0.01). Importantly, when we included the non-brain traits in this analysis, we saw on average minimal number of significantly enriched pathways (mean = 1) showing that our method correctly prioritizes the brain-related traits in a brain single cell atlas.

Finally, we investigated if there are any cell type specific regional effects in brain trait association. Using the same strategy as the MDD cell type region analysis, we found few region cell type pairs with high specificity index (specificity index > 2) (**Supplementary Figure 17 and 18**). The maxima for number of associations with high specificity index are observed in L6-IT neurons at Brodmann region 19 for intelligence savageness, verbal numeric reasoning, and BMI. Most of the cell type region pairs that do have a high specificity index are unique to one trait. For example, social well-being has unique association with L5-IT neurons, where primary motor cortex, prefrontal cortex, inferior temporal gyrus, Brodmann area 19 and Brodmann area 46 only have a high specificity index in the L5-IT context for social well-being trait. Taken together, we reason that although there are cell types such as L2/3-IT neurons that is broadly associated with brain traits, there are still differences in risk association along spatial axis in a cell type specific manner.

### Met-scDRS for CpG methylation

Interested in how met-scDRS can be applied in CpG methylation context, we also applied met-scDRS on human methylome atlas dataset as well (**Methods**) and visualized the significance in associations across traits and cell types (**Figure 6c, d**, **Supplementary Figure 19, Supplementary Table 6**). Across 75 tested traits, most of the significant (FDR corrected p value < 0.1) associations are in brain related traits (**Supplementary Figure 19, Supplementary Table 6**). Like our observation where L2/3-IT excitatory neurons are strongly associated with MDD (**Figure 6b, c, d**), we observed 36.3% of all L2/3-IT excitatory neurons are significantly associated with MDD in CpG methylation. This abundance in significant associations is followed by L5-ET neurons where we observe 31.1% of all L5-ET neurons are significantly associated with MDD. Another similarity between CpG and non-CpG methylation results is that MSN inhibitory neurons expressing D2 receptors were also well associated with SCZ in CpG methylation. These observation shows us Met-scDRS can identify common disease associations between CpG and non-CpG methylation. However, there are also dissimilarities between the two modalities (**Figure 6b, d**). One such example is dentate gyrus (DG) neurons’ association with bipolar disorder. In CpG methylation, 66.0% of the DG neurons are associated with the disease, where its association was minimal in non-CpG methylation. Another of such example is in Foxp2 inhibitory neurons, where in non-CpG methylation, it has limited association to brain related traits (**Supplementary Table 2**), in contrast to the moderate association in SCZ (> 14% in proportion) in CpG methylation (**Supplementary Table 6**). This result demonstrated met-scDRS is portable to CpG methylation and obtains disease associations complementary in the two methylation profiles.

### Met-scDRS captures disease association beyond transcriptomics

Finally, we tested how our findings in met-scDRS on MDD can be replicated in transcriptomics analysis. Using two human transcriptomics datasets, one with case/control transcriptomics^39^, and another an atlas level transcriptomics dataset^40^ released by Allen Institute for Brain Science (AIBS); we applied scDRS with MDD gene set to identify disease implicated cells. Interestingly, we did not find any significant cells that pass multiple testing correction (FDR < 0.1). However, using only the set of cells that are marginally significant (p value < 0.05) in both transcriptomic datasets, we still identified the excitatory neurons showing strong enrichment in signal as our methylation atlas analysis (**Supplementary Figure 20a, b**). It is also worth pointing out that the number of inhibitory neurons in both datasets show slightly higher number of marginal significance than excitatory neurons (**Supplementary Figure 20a, b**). Importantly, excitatory neurons at L2/3 still shows strong marginal significance in both datasets (**Supplementary Figure 21a, b**). Specifically, in atlas data, 403 excitatory neurons from layer 2 and 3 are marginally significant, representing 12.38% of all marginally significant cells while in case/control transcriptomics dataset, we identified 639 marginally significant L2-4 excitatory neurons, representing 20% of all marginally significant cells. Non-neuronal cells continue to represent the least amount of marginal significance, with only 11.1% and 6.4% of the marginal significance in case/control dataset and AIBS transcriptomics dataset respectively. Although the transcriptomics analysis suggests a similar trend, we clearly observe a much higher number of significant cells after FDR correction in MDD from methylomics. We reason that met-scDRS can provide additional level of information beyond transcriptomics and to allow researchers identify new cell types that are associated with diseases.

## Discussion

We present met-scDRS as a statistical tool that finds polygenic disease associations at single cell resolution and prioritizes disease implicated cells and genes. Using simulation studies, we show that our method is well powered and calibrated under a variety of different scenarios. When applied to human methylation atlas level data using MDD GWAS genes, we identified glutamatergic neurons as the main associated neuronal class with substantial cell type heterogeneity within the neuronal class. This result showcases the power of met-scDRS to utilize non-CpG and CpG methylation, which is highly cell type specific and involved in cell type establishment^21,22,24,41–43^. Met-scDRS shows inhibitory neurons such as MSN neurons expressing D2 receptors is highly significant in disease association in SCZ in both of the methylation modalities. This is note-worthy for two reasons, first there is strong existing evidence of the importance of MSN in SCZ^44–48^, and two, D2 receptors are one of the main druggable targets by existing antipsychotics^49–52^.

Met-scDRS computes disease risk association at a single cell resolution thus being able to identity heterogeneity across cells. For example, within the temporal gyrus, a known region with MDD associations^53,54^, met-scDRS identifies the association gradient in L2/3-IT neurons from middle temporal gyrus to inferior. Met-scDRS can also be utilized as a gene prioritization tool where we recovered synapse related pathways are primarily enriched in MDD compared to the GWAS genes, which suggests that hypomethylation in these genes could be risk conferring and should be further researched. This result coincides and supports the current understanding of dysfunctional neurotransmitter and synaptic pathways dysregulation in MDD^29–33^. Since non-brain traits usually do not involve brain tissue while brain related traits do^55,56^,we consider the lack of signal in non-brain related traits to suggests that met-scDRS is well calibrated in real world scenarios. Another important feature is the portability between CpG and non-CpG modalities by met-scDRS. Since CpG and non-CpG differ in target methylation cell types^41^, having the capacity to infer disease association in both profiles allow researchers to 1) utilize under-researched non-CpG methylation modality^57,58^ and 2) gauge functional convergence and divergence in the respective sequencing profile. For example, we were able to identify L2/3-IT neurons as a major source of disease association in both CpG and non-CpG methylation for MDD while DG neurons, a critical neuronal type for learning and memories^59^, is only strongly associated with Bipolar and educational traits in CpG methylation.

We conclude with a few limitations in our approach. First, met-scDRS could underestimate the contribution of disease association when the cell has low global methylation levels. For example, non-CpG methylation is lowly expressed in non-neuronal cells (**Supplementary Figure 5-7**), which could explain the lack of significant associations. Thus, researchers should be mindful of such false negatives for cell types with known minimal expression of methylation. We refer to other tools and sequencing modalities for computing true disease association for these cases. Since CG sites are much sparser in CA sites in the genome^34^, we used K-nearest neighbor aggregation to do reliable inference when utilizing met-scDRS for CpG methylation. Although it accounts for genomic sparsity, such aggregation reduces resolution and could potentially introduce bias. Second, there is the possibility where there is a strong pleiotropic effect with traits, such as atrial fibrillation, met-scDRS can show abundant and significant associations within a dataset sequenced from brain cells. Although atrial fibrillation is classified as a heart related trait, it has strong neuronal basis where the mechanism of the disease could in part be attributed to the excitability of the atrial cardiomyocytes^60,61^. Such pleiotropic effect could manifest in met-scDRS as significant associations since the genes associated with atrial fibrillation with respect to such excitability^61^ could also be particularly perturbed in methylation in brain region. This demonstrates that false positives could arise with pleiotropic effect. Third, there may be false positive genes in the putative disease gene set aggregated over GWAS using tools such as MAGMA^62^. Although we have shown that the power of met-scDRS increases linearly with the percentage of true positive genes in the putative gene set in limitation, it could still lead to a decrease in power. Considering this, we still chose GWAS genes summarized by MAGMA as a data driven approach in probing how the disease associated variants detected in GWAS can manifest in hypomethylation context. Met-scDRS could also circumvent this limitation by taking in a user specified gene set such as SFARI^63^ gene set for autism disorder where putative disease genes are meticulously curated and ranked by publication related evidence. Such approach can provide disease mechanistic insights and allow researchers to understand if their curated gene set using techniques such as forward genetic screening could have an association with diseases via DNA methylation modification. Finally, it is also worth noting that some of the dissection regions are not present in the human brain region atlas released by AIBS, while other regions are too course as they contain multiple dissection regions in the human methylation atlas dataset.

## Methods

### Single cell dataset acquisition

We utilized two single cell methylomics datasets, where one of them contains human methylome, and one contains mouse methylome under GEO ascension number GSE215353 and GSE132489. In GSE132489, single nucleus methylation cytosine sequencing (snmC-seq) of 92 adult mouse (mouse methylation brain atlas) was downloaded from gene expression omnibus GSE132489 on 08/17/2023. This dataset contains 103,982 methylomes that passed QC across 17 dissection regions with 41 major cell types profiled by snmC-seq2. In GSE215353, we downloaded 373,888 single cell DNA methylomes across three adult brains spanning 44 regions profiled by snmC-seq2 (human methylation brain atlas) containing 40 major cell types on March 22, 2024. Additionally, we also analyzed two transcriptomics datasets, one with case/control transcriptomics^39^, and another a atlas level transcriptomics dataset^40^ released by AIBS. The case/control transcriptomics dataset is downloaded in May 2023 while the atlas level transcriptomics dataset is downloaded in December 2024.

### 75 traits magma gene set acquisition

For the GWAS datasets, we first downloaded the same gene set files derived from 74 UK BioBank traits as in the previously published work^19^ for met-scDRS. Then, we computed the MAGMA gene set using autism disorder GWAS^64^ and combined the resulting gene set with the original 74 traits. To describe the processing scheme of Autism disorder GWAS in detail, we first downloaded the gene location file and SNP location file from MAGMA^62^ website https://ctg.cncr.nl/software/magma for GRCH37 genome build. We then filtered the SNPs in the autism disorder GWAS to the set of Hapmap3^65^ SNPs. Next, using the gene and SNP location files that we downloaded, we used MAGMA to generate a gene annotation file with 10kb as our window size both upstream and downstream of the gene location. Finally, using the filtered autism disorder GWAS, we computed the MAGMA genes using the p values of the SNPs within the GWAS and the total number of individuals participated in the GWAS study. The top 1000 genes from the resulting MAGMA gene set file with their respective Z scores are extracted and aggregated to the 74 UK BioBank traits gene set.

### met-scDRS Algorithm

For the methylationCytosine sequencing dataset, met-scDRS takes gene level non-CpG methylation ratio as input where the ratio is defined as methylcytosine base coverage divided by the total cytosine base coverage aggregated at gene bodies. The purpose of such aggregation at gene body substantially decreases the data dimension and sparsity. Met-scDRS calculates the inverse fraction of methylated cytosine *X*’*_c,g_* as 1 - *X*’*_c,g_* where x is the aggregated ratio at cell *c* and gene *g* as methylation level are expected to be anticorrelated to the gene expression.

It should be noted that such expectation albeit supported by known genetic regulation mechanism^41^, is not always observed^23^. We then filter off genes that have very low variance (< 5th percentile) for variance stabilization. For CpG methylation modality, due to the sparsity in CG base in the genome^34^, we take an additional step to aggregate the inversed and variance filtered *X’_c,g_* for each cell using 5 nearest neighbors in Euclidian distance at UMAP space. Finally, we use the existing algorithm detailed in scDRS^19^ for obtaining cell-wise z score (met-scDRS) and p values. In brief, the procedure would partition genes based on average inverse fraction and variance and select background genes with matching variance level and average inverse fraction level for computing a null distribution. For further details, we refer to the original scDRS publication^19^, as well as our publicly available code implementation (https://github.com/bogdanlab/met-scDRS).

### Calibration simulations

From the GSE132489 mouse methylation brain atlas dataset, we first filtered to the set of cells that pass QC detailed in the publication^21^ and randomly selected 10,000 cells. For these 10,000 cells, we computed *X’_c,g_* and filtered for set of genes with > 5th percentile variance as our simulation methylation matrix. From this matrix, we further randomly selected three sets of genes: 1) genes randomly selected from all the features in the simulation matrix. 2) Top 25% of genes with the highest fraction. 3) Top 25% of genes with the highest fraction variance. To account for the effect of gene sizes during simulation, sets of 100 genes, 500 genes, and 1,000 genes are selected. Next, we permuted the GWAS weights by randomly selecting one trait out of the downloaded 74 traits gene set file and then randomly selecting the top 100, 500, and 1,000 z scores respectively to pair with the simulated gene set. We also considered a more stringent threshold for simulating the high mean and variance of genes where genes are selected from 1) top 5% of genes with the highest fraction 2) top 5% of genes with the highest fraction variance using the same procedure mentioned above. We repeated all simulations with 100 replicates to estimate the standard deviation between simulations. In total, we have 100 replications x 3 gene sets sizes x 6 simulations for calibration simulation.

### Causal simulations

From X’*_c,g_* obtained in the previous null simulation, we perturbed the methylation values in two different ways: 1) fixed overlap and 2) fixed effect size. For fixed overlap, we first select 500 cells and 1,000 genes from *X’_c,g_* randomly as causal genes and causal cells. Out of these 1,000 genes, only 250 (fixed 25% overlap) genes are chosen from human ortholog genes present in the gene set file of height trait. The remaining 750 genes are then chosen randomly from all genes present in *X’_c,g_*. For these 500 cells and 1,000 genes, their values in the *X’_c,g_* are multiplied by 1.01 to 1.05 at 0.01 increments as well as from 1.001 to 1.01 at 0.001 increments to observe met-scDRS behavior across a range of perturbations. The perturbed x1 is capped at 1 such that all values after perturbation remain bound between 0 and 1 to retain the processed data in the fraction space. For fixed effect size casual simulation, we vary the overlap previously held at 25% while keeping the perturbed effect size constant at 1.005 times the original *X’_c,g_*. We explored a range of overlaps from 10% to 50% at 10% increments. For causal simulation, the gene weight from height trait is used. We repeated all simulation settings for 100 times and quantified power as percentage of perturbed cells correctly identified as significant after FDR correction (< 0.1) by met-scDRS.

### met-scDRS application on GSE215353

For GSE215353^22^, since the downloaded dataset is already preprocessed, we did not deploy additional preprocessing and inherited the cell type labels from the original publication’s metadata for downstream analysis. For CpG methylation modality, due to the sparsity in CG base in the genome^34^, we aggregated CpG methylation for each cell using 5 nearest neighbors in Euclidian distance at UMAP space. Then for both of the modalities, we applied the met-scDRS to the dataset using the 75 traits GWAS gene set file for both CpG and non-CpG methylation profiles. We obtained the cell wise met-scDRS disease z score and p values for all traits and corrected the p value using FDR correction within each of the traits and quantified the number of significant cells (< 0.1 FDR corrected p value). We visualized the proportion of significant cells in each of the cell type. We also visualized the met-scDRS in UMAP space using the original publication provided coordinates. On non-CpG methylation, we calculated the z score average for each cell type in each region for each of the 75 traits for visualization and tested for cell type specific dissection region heterogeneity to investigate if there is a region-specific cell type effect. On a separate analysis, we also included sequencing pool, batch, donor and scaled methylation expression as covariates and regressed them out to ensure that our observations are not confounded by batch effects. To reduce computational burden, we only computed batch effect corrected analysis and CpG methylation modality analysis on a random subset of data of 74,777 cells while the non-CpG methylation is computed on the full dataset.

### GSE215353 region rendering

Using the met-scDRS average at each cell type in each region, we mapped the dissection regions released in GSE215353 onto the human adult brain reference frame (version 0.1) released by Allen Institute for Brain Science^66^ (AIBS), summarized in **Supplementary Table 3**. It should be noted that there are some ambiguities in the mapping since some of the dissection regions are not present in the human brain region atlas released by (AIBS atlas) and some brain region in the AIBS atlas is too coarse that it contains multiple dissection regions released in the GSE215353 data. For regions in the reference frame with multiple mapping of dissection regions from the GSE215353 dataset, we aggregated the entries by taking the average of the multiple mappings and dropped the regions that does not exist in the reference frame or outside of the common cortex axis. The resulting average MDD met-scDRS at each region are used as color scale for the visualization and rendered with brainrender^67^ using python v3.10.10.

### Cell type specific dissection region heterogeneity testing

To test if a region is significantly associated with disease risk score in a cell type specific manner, we developed a cell type region specificity index using the following algorithm: firstly, for each of the 75 traits, we subset the set of cells that have FDR corrected p value < 0.1. Secondly, we subset to the cells within each of the cell type. Thirdly, for each of the dissection region in the cell type that has more than 100 cells, we build a linear model by coding a binary covariate label on if the cell originates from this region and estimate the effect size of region on predicting the normalized score computed using both putative disease genes and control genes. The constraint on number of cells is to ensure robustness while the utilization of both normalized score for both putative disease genes and control gene set is used to mitigate the *i.i.d* assumption

Mathematically with dissection region *j* as an example:

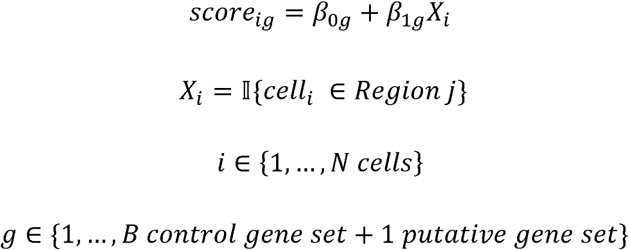

Since the control genes are generated via Monte Carlo sampling on genes that matches the methylation level and the variance of the putative disease gene set, we compute the specificity index as how far away *β*_1_ is in putative gene set compared to those derived from the *B* control gene set. Mathematically:

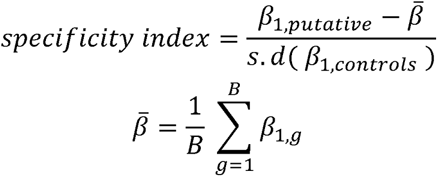

Iterating through all regions, cell types, and 75 traits, we compute a tissue specificity score for each cell type, tissue, trait triplet with more than 100 significant cells.

### Gene ontology analysis

For each gene, we computed the spearman correlation between methylation level of *X’_g_* and met-scDRS for all 75 traits. Then, for each of the 75 traits, we selected the top 100 genes with the highest spearman correlation coefficients as our foreground gene sets. We interpret each spearman correlation as each gene’s met-scDRS contribution and provide a mean to quantify how disease relevant each gene is. For the background gene set, we used the set of 1000 GWAS MAGMA genes present in *X*’, i.e.: we used the disease gene set input for met-scDRS as a way to control for background putative genes that is already known to be associated with disease. Using this approach, we can know what genes are important for disease association on top of set of MAGMA genes which is known to broadly associated with the trait *a priori*. Using the selected foreground and background set, we tested for pathway enrichment within the C5 ontology released by GSEA^68–70^ with ClusterProfiler^71^. For each of the 75 traits, we visualized the top 10 over-represented gene ontologies and the genes from the foreground that are the most associated with the ontologies in a network diagram.

### Transcriptional analysis

We downloaded two human transcriptomics datasets, one with case/control transcriptomics^39^, and an atlas level transcriptomics dataset^40^ released by AIBS. For the case/control MDD dataset, we corrected for batch effect by including the batch variable as a covariate in the scDRS pipeline. For the AIBS transcriptomics dataset, we corrected for donor and sex effect. We supplied the count matrix as input for scDRS to obtain risk score estimates for MDD gene set. Since no cells remain significant after FDR correction, we visualized the count of marginally significant cells among cell types and cell class at different layers.

