## Supplementary figures for "Integration of polygenic risk with single cell methylation profiles for depression"

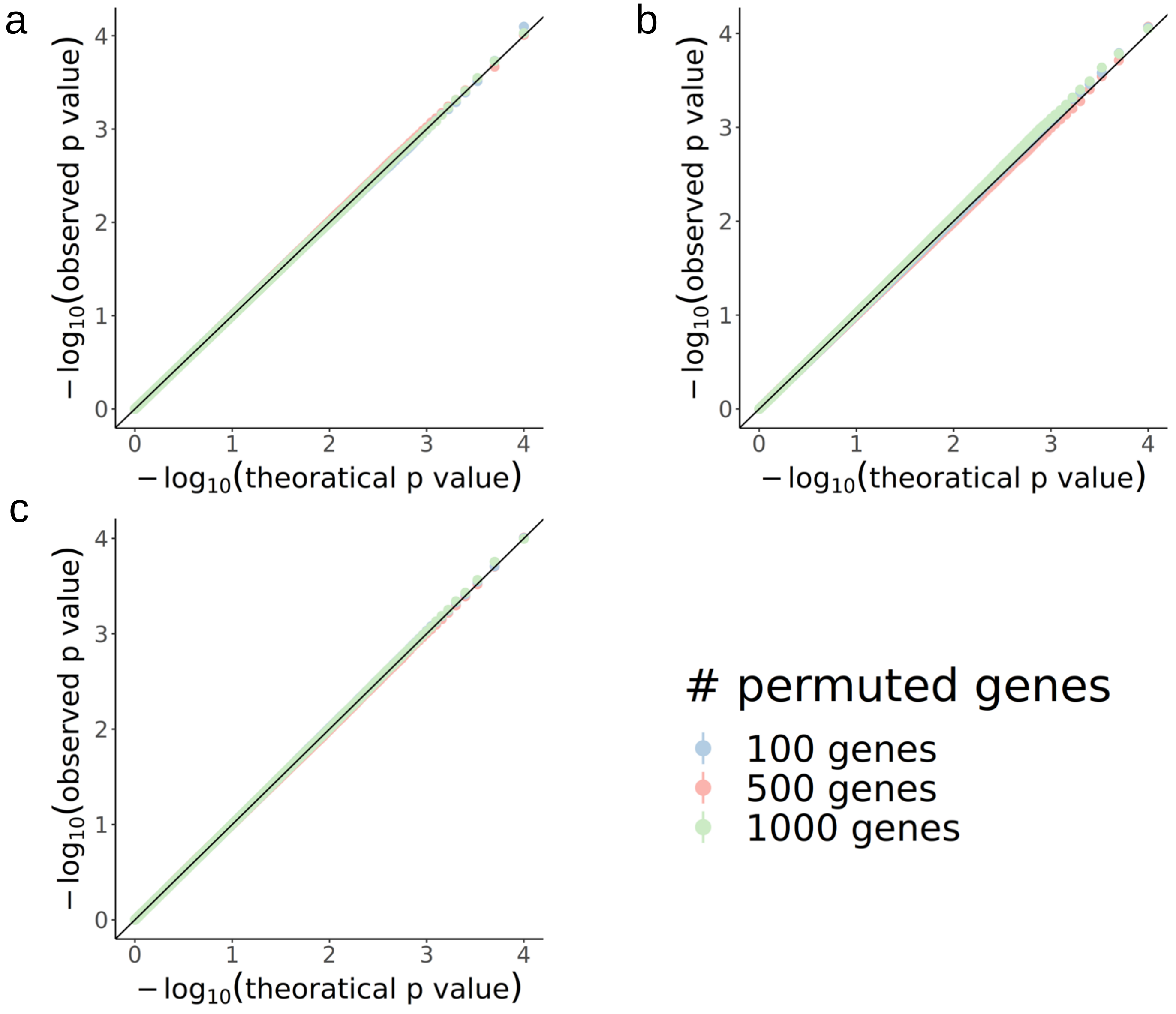


**Supplementary figure 1: Calibration curve corresponding to genes draw (a) randomly, genes drew from genes with (b) top 95$^{th}$ percentile in variance and (c) 95$^{th}$ percentile in inverse fraction$(X'_{c,g})$. All calibration curves are colored by number of permuted genes within the simulated gene sets. Met-scDRS is run with gene weight option set to variance stabilization weights**

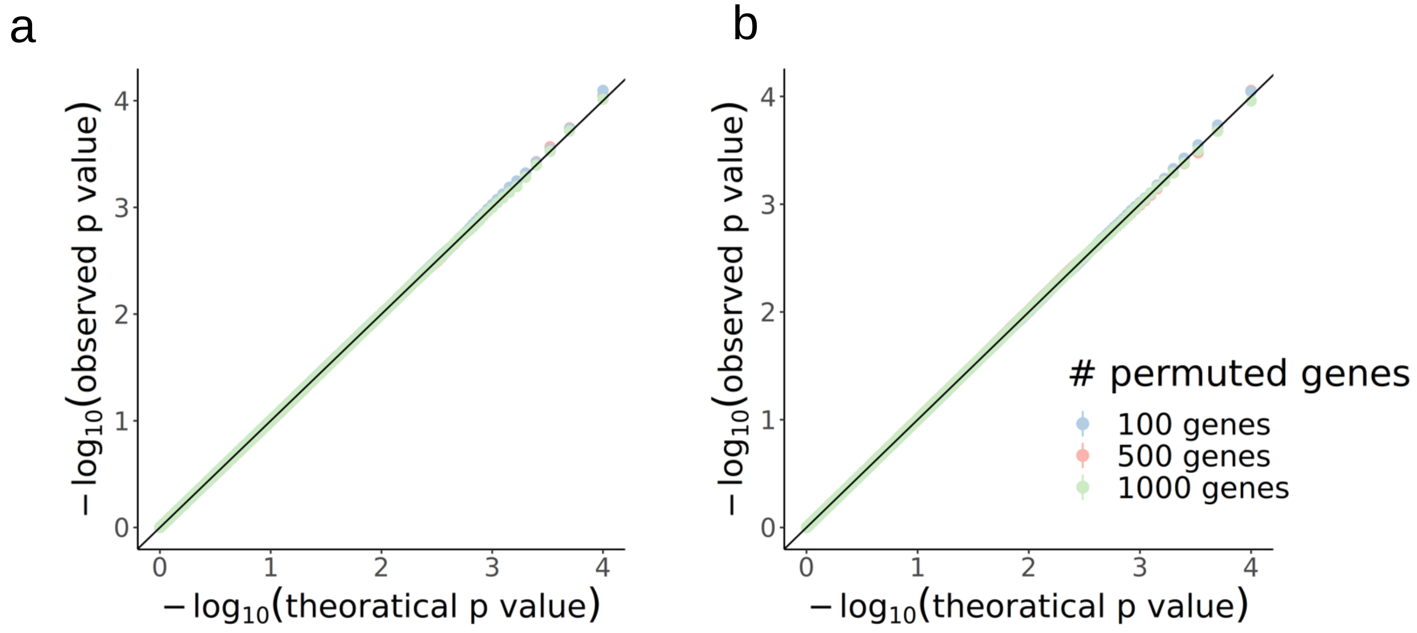


**Supplementary figure 2: Calibration curve corresponding to genes draw from (a) top 75th percentile in variance and (b) 75th percentile in inverse fraction$(X'_{c,g})$. All calibration curves are colored by number of permuted genes within the simulated gene sets. Met-scDRS is run with gene weight option set to variance stabilization weights**

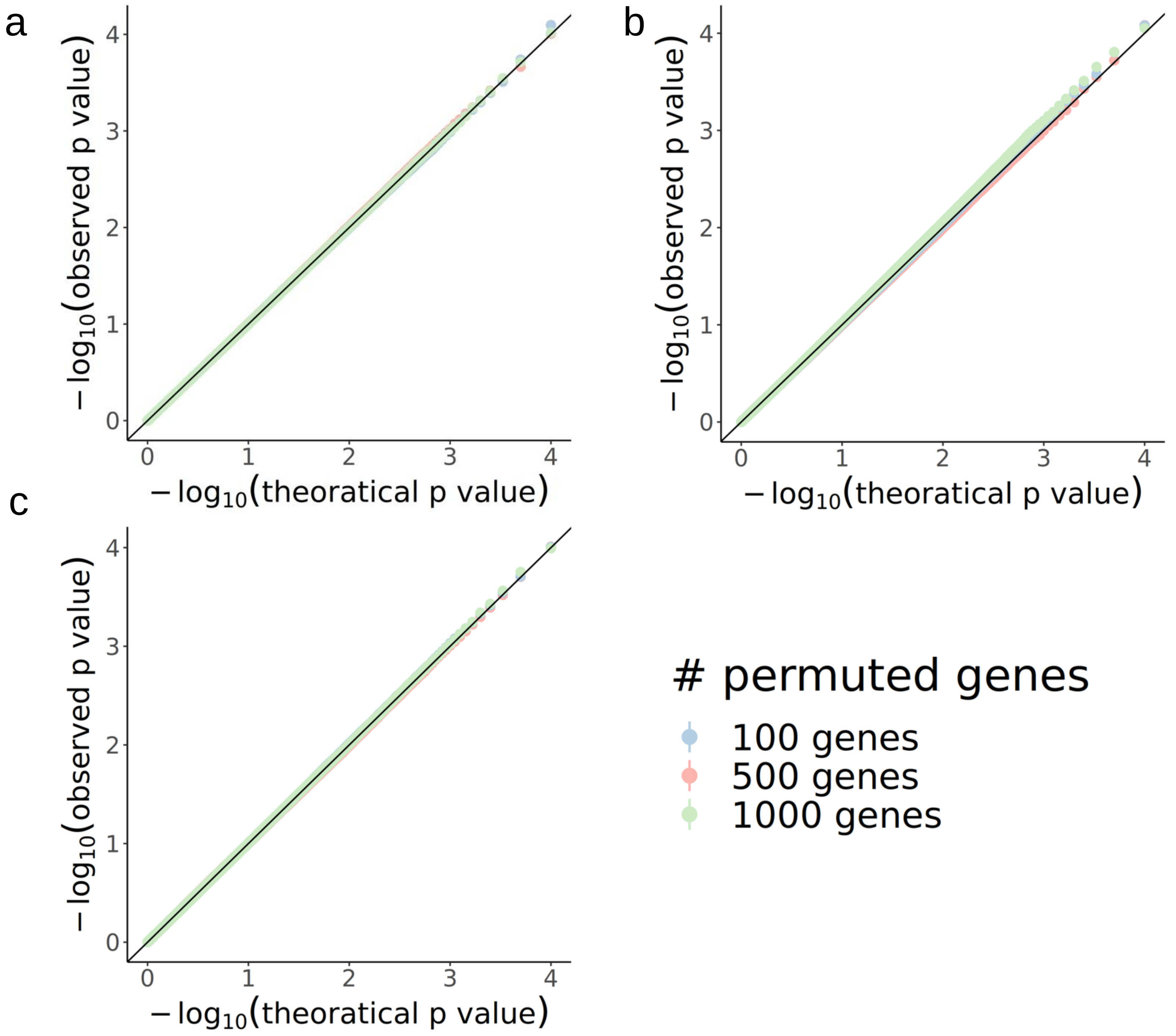


**Supplementary figure 3: Calibration curve corresponding to genes draw (a) randomly, genes drew from genes with (b) top 95th percentile in variance and (c) 95th percentile in inverse fraction$(X'_{c,g})$. . All calibration curves are colored by number of permuted genes within the simulated gene sets. Met-scDRS is run with gene weight option set to inverse standard deviation**

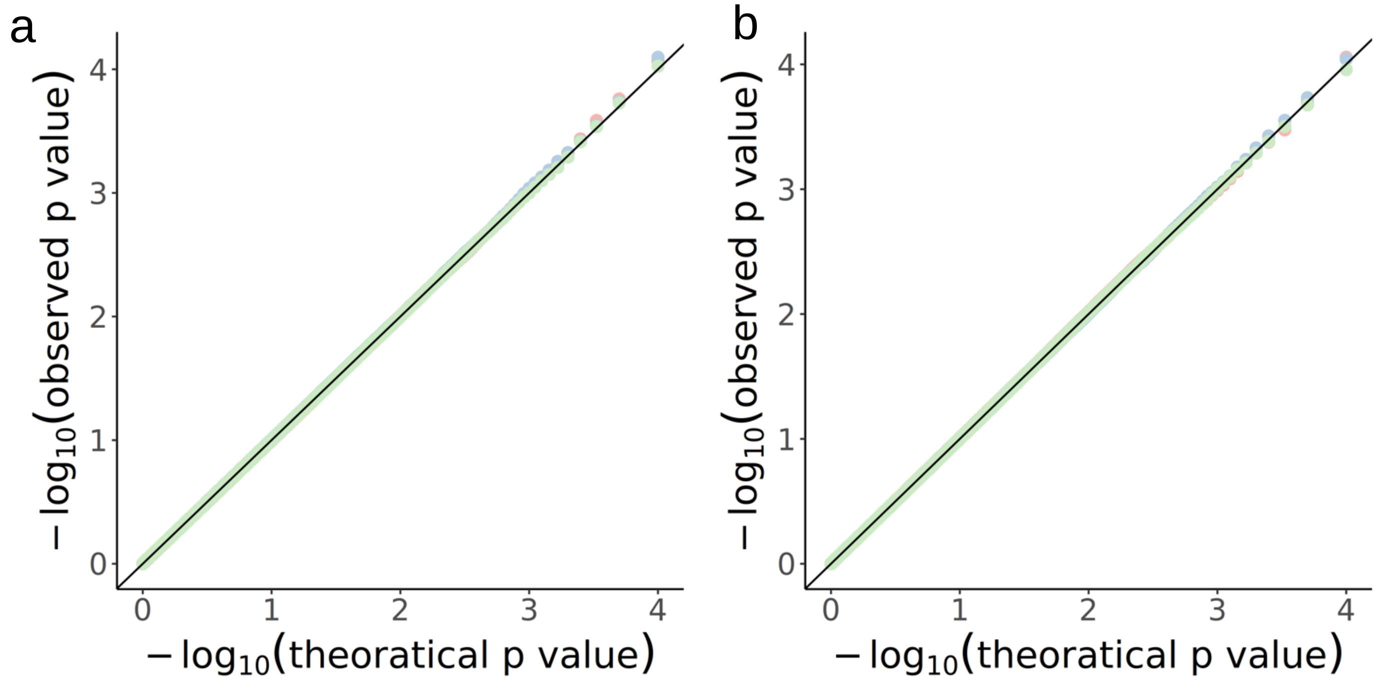


**Supplementary figure 4: calibration curve corresponding to genes draw from a) top 75$^{th}$ percentile in variance and b) 75$^{th}$ percentile in inverse fraction$(X'_{c,g})$. All calibration curves are colored by number of permuted genes within the simulated gene sets. Met-scDRS is run with gene weight option set to inverse standard deviation weights**

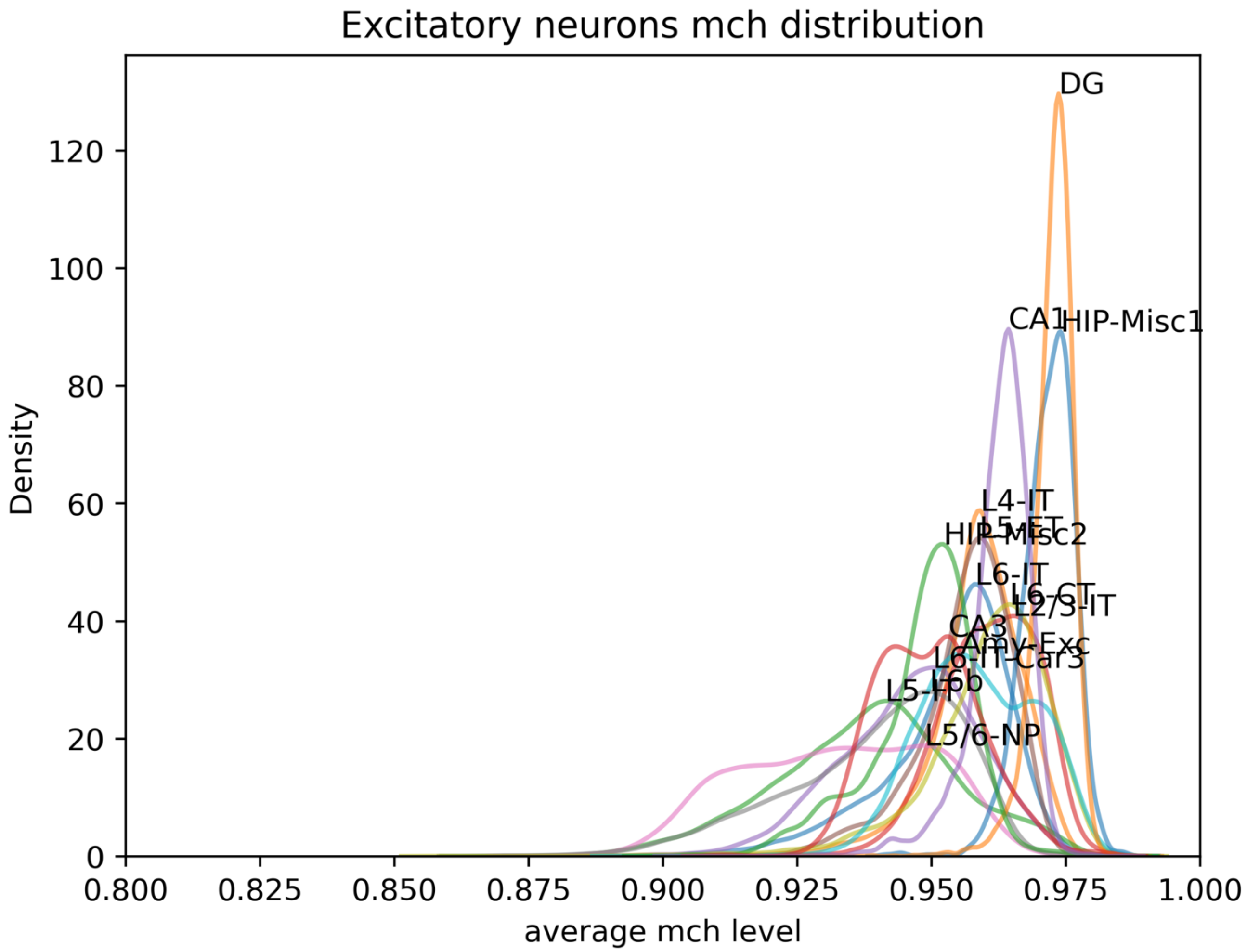


**Supplementary figure 5: Density plot of average methylation level in $X'_{c,g}$. Colored by each cell type in Excitatory neuronal class. Cell type is directly labelled at the peak of the distribution**

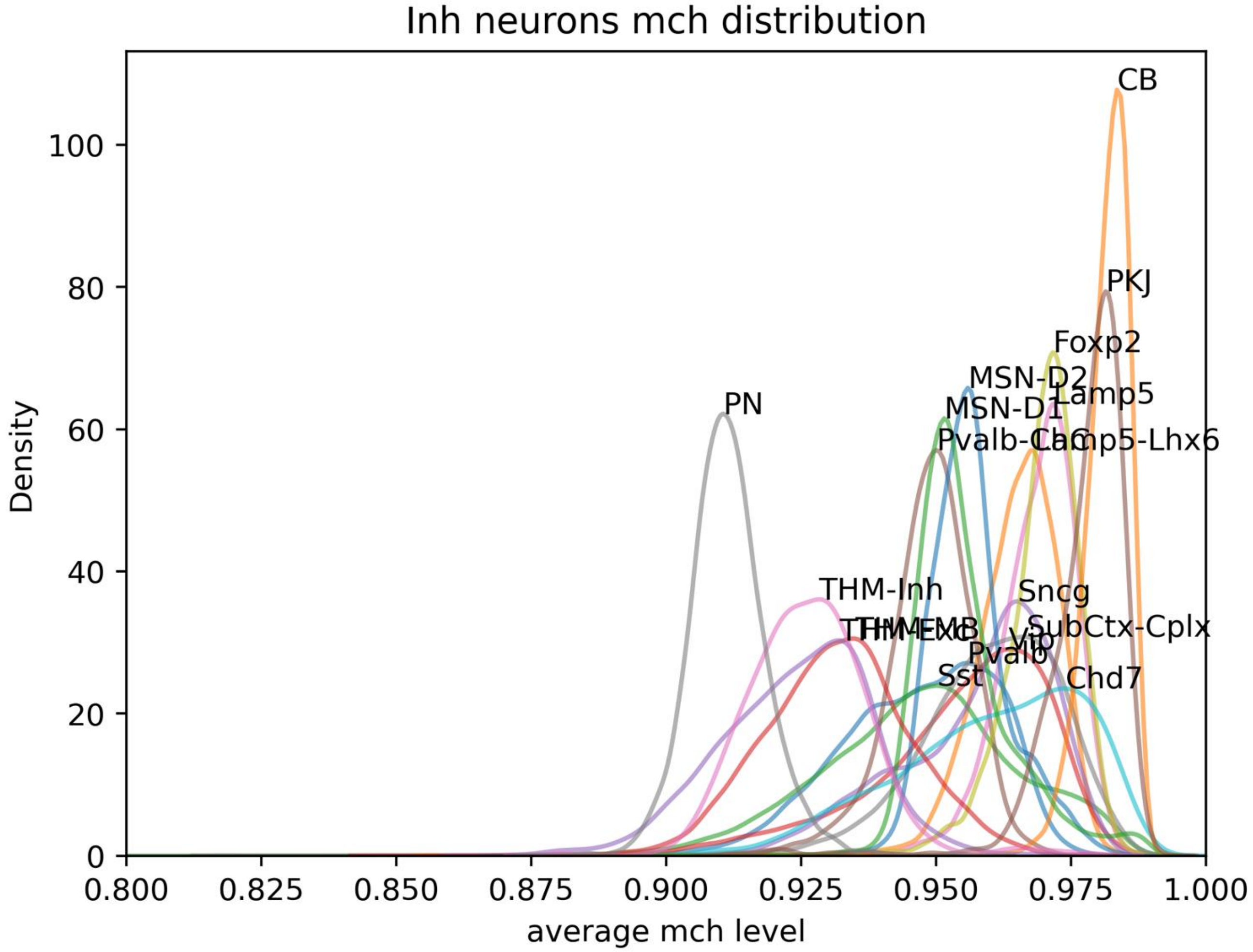


**Supplementary figure 6: Density plot of average methylation level in $X'_{c,g}$ colored by each cell type in Inhibitory neuronal class. Cell type is directly labelled at the peak of the distribution**

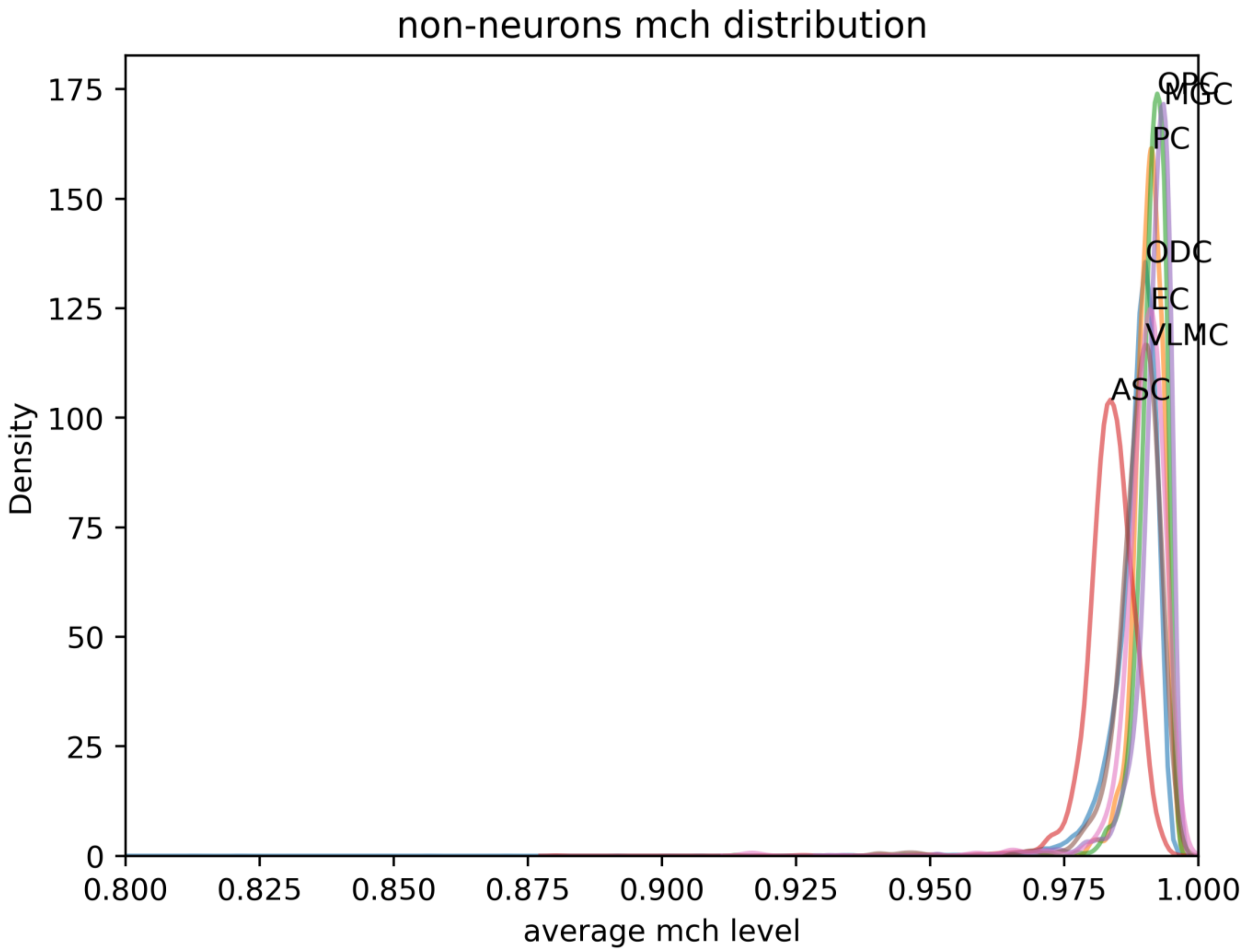


**Supplementary figure 7: Density plot of average methylation level in $X'_{c,g}$ colored by each cell type in non neuronal class. Cell type is directly labelled at the peak of the distribution**

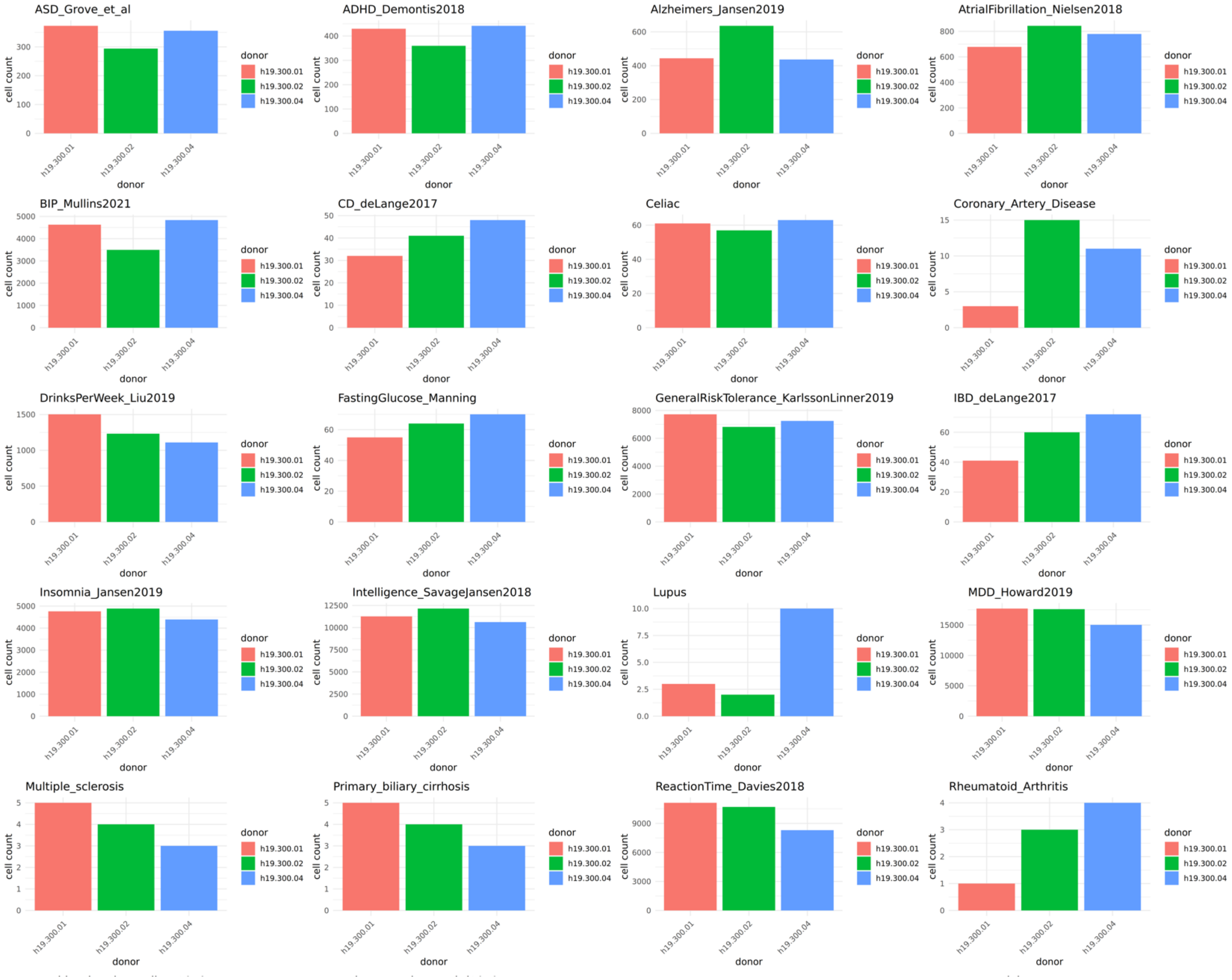


### Supplementary figure 8: Number of significant cells colored by donors in each disease

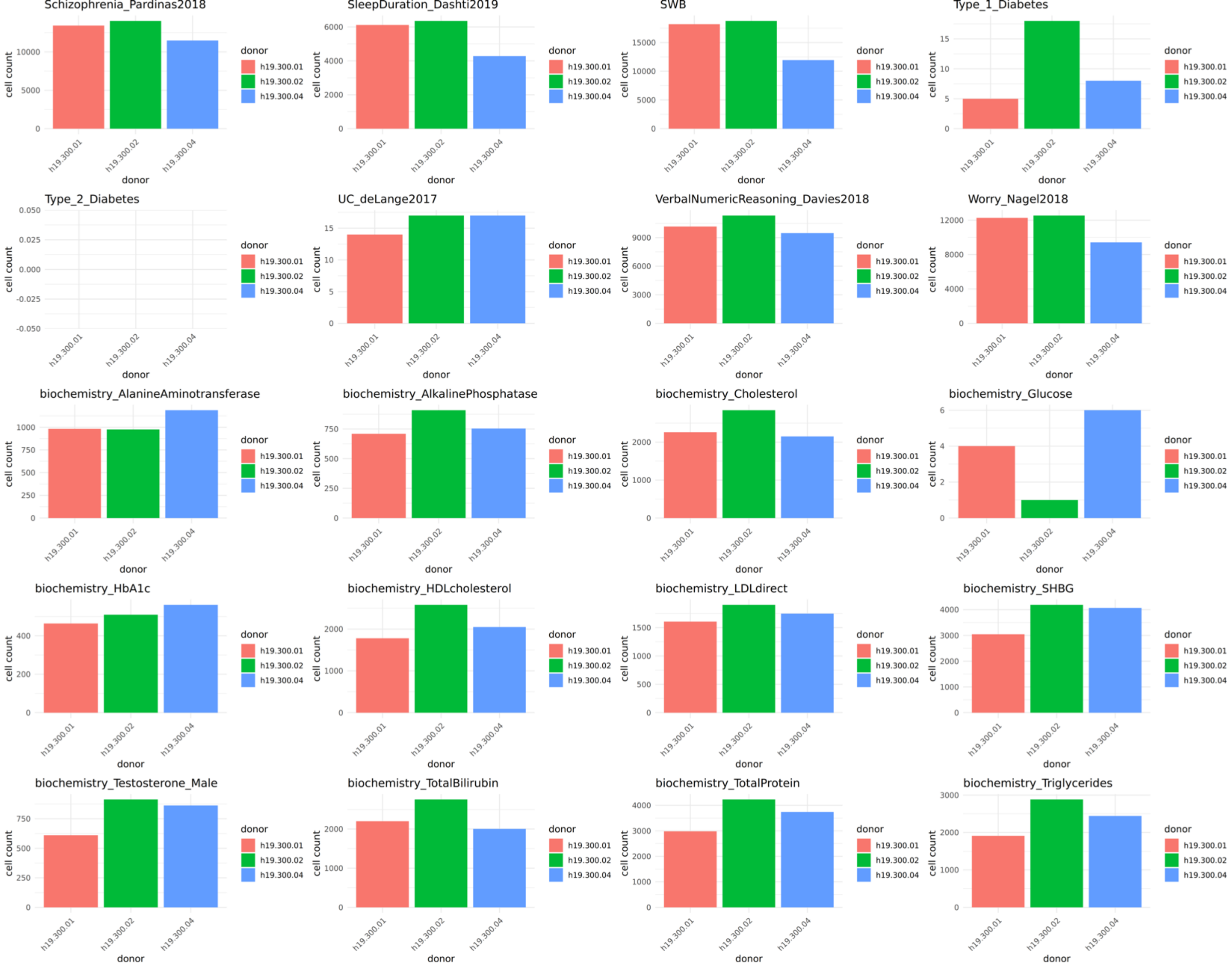


**Supplementary figure 8: Number of significant cells colored by donors in each disease (continue)**

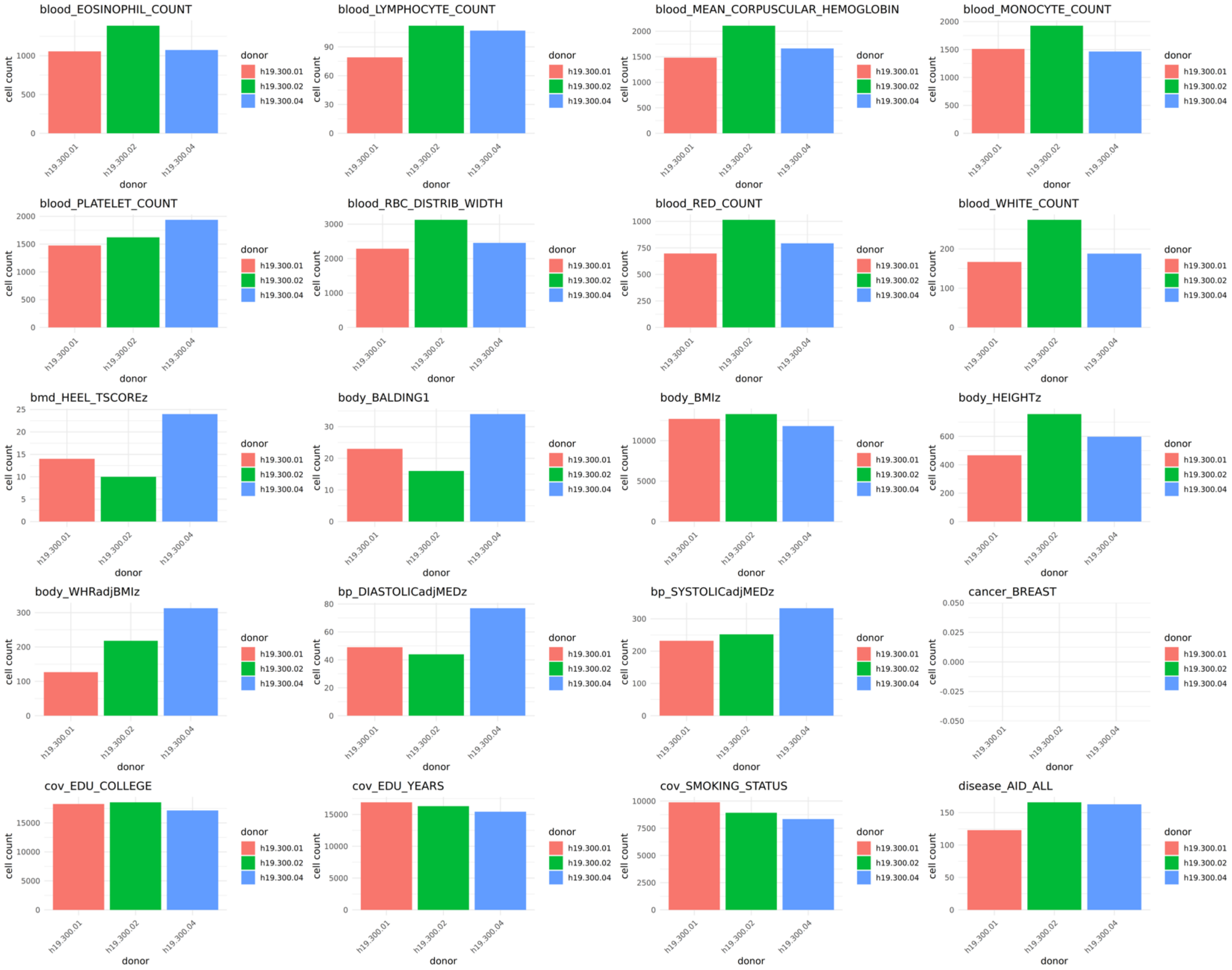


### Supplementary figure 8: Number of significant cells colored by donors in each disease (continue)

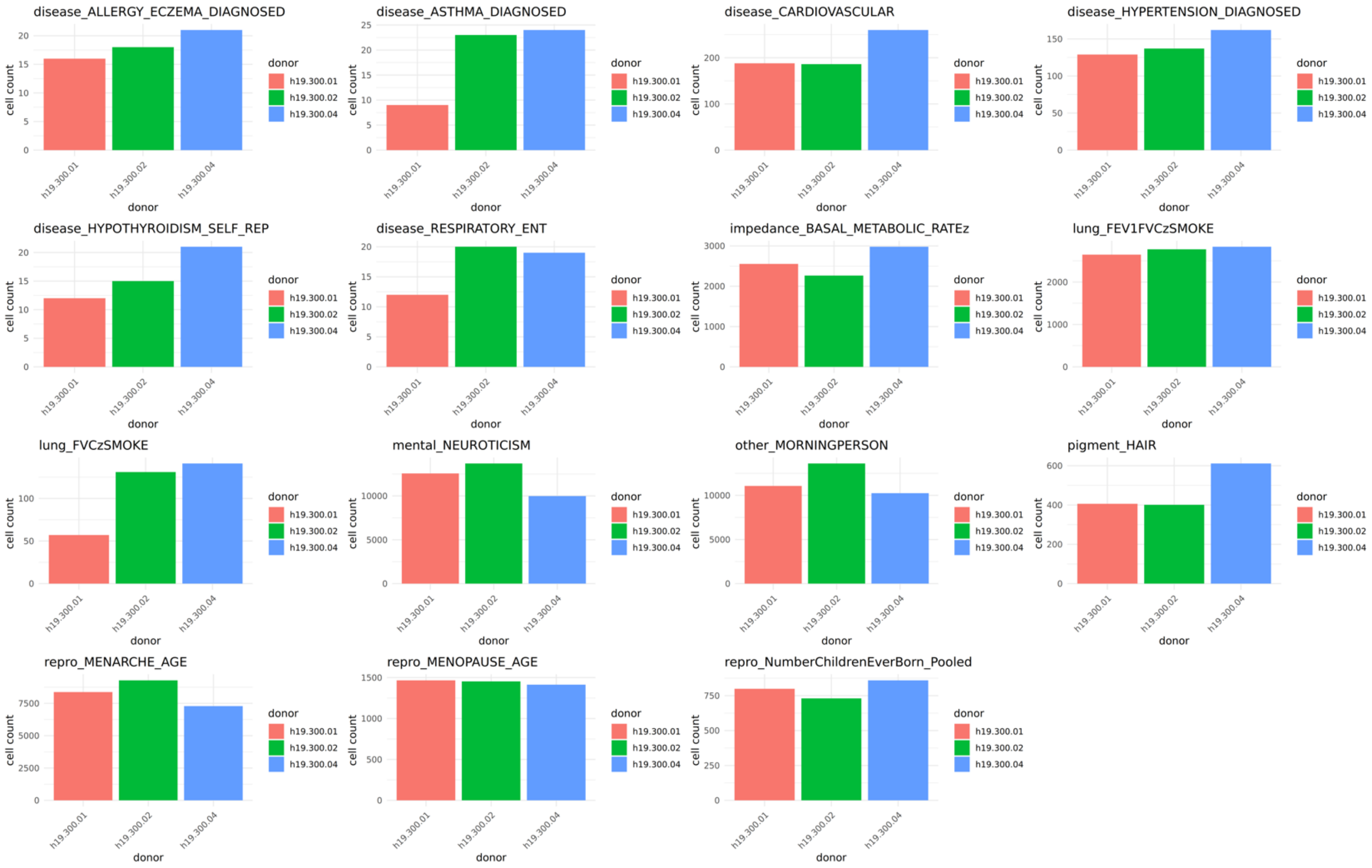


**Supplementary figure 8: Number of significant cells colored by donors in each disease (continue)**

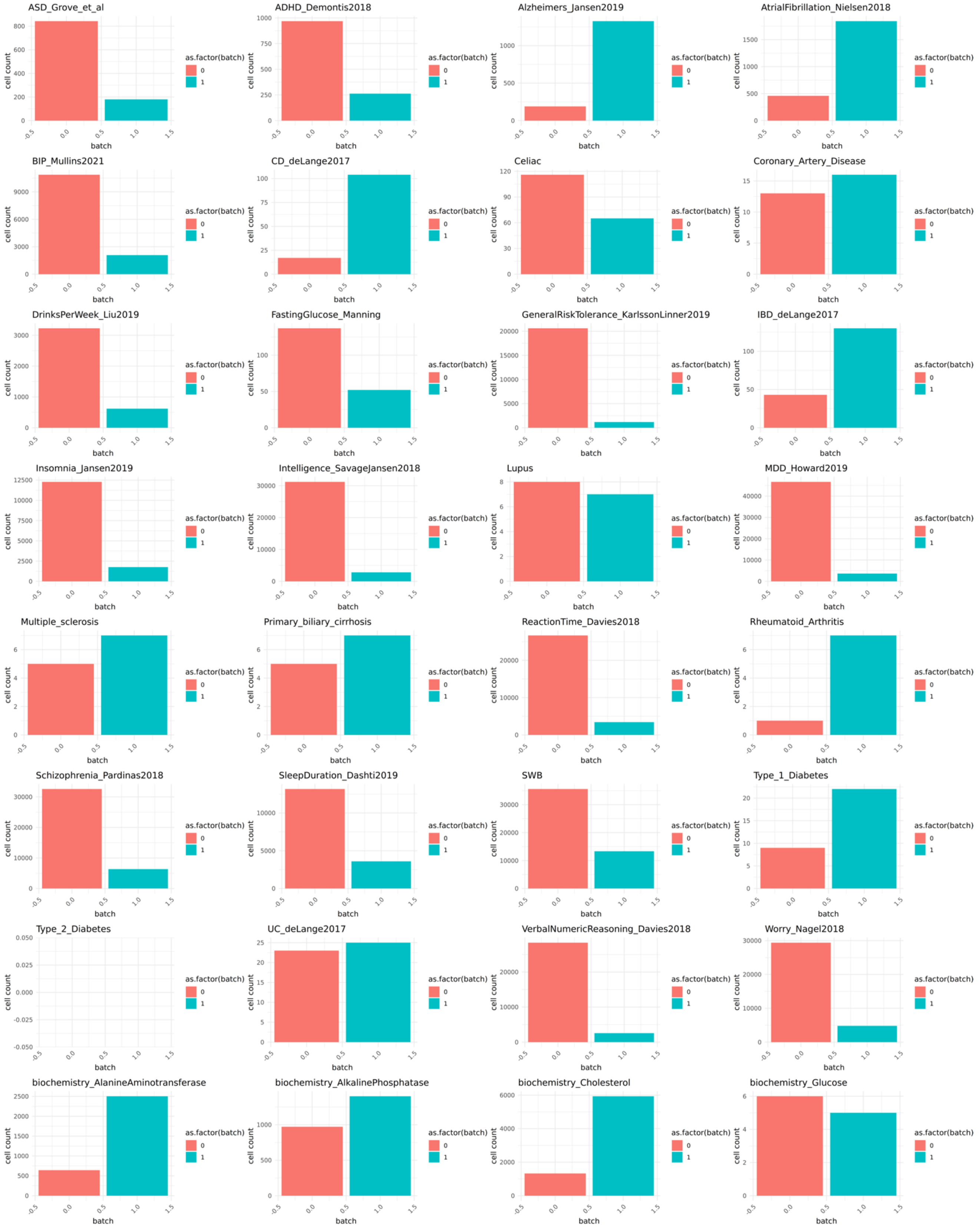


### Supplementary figure 9: Number of significant cells colored by batch

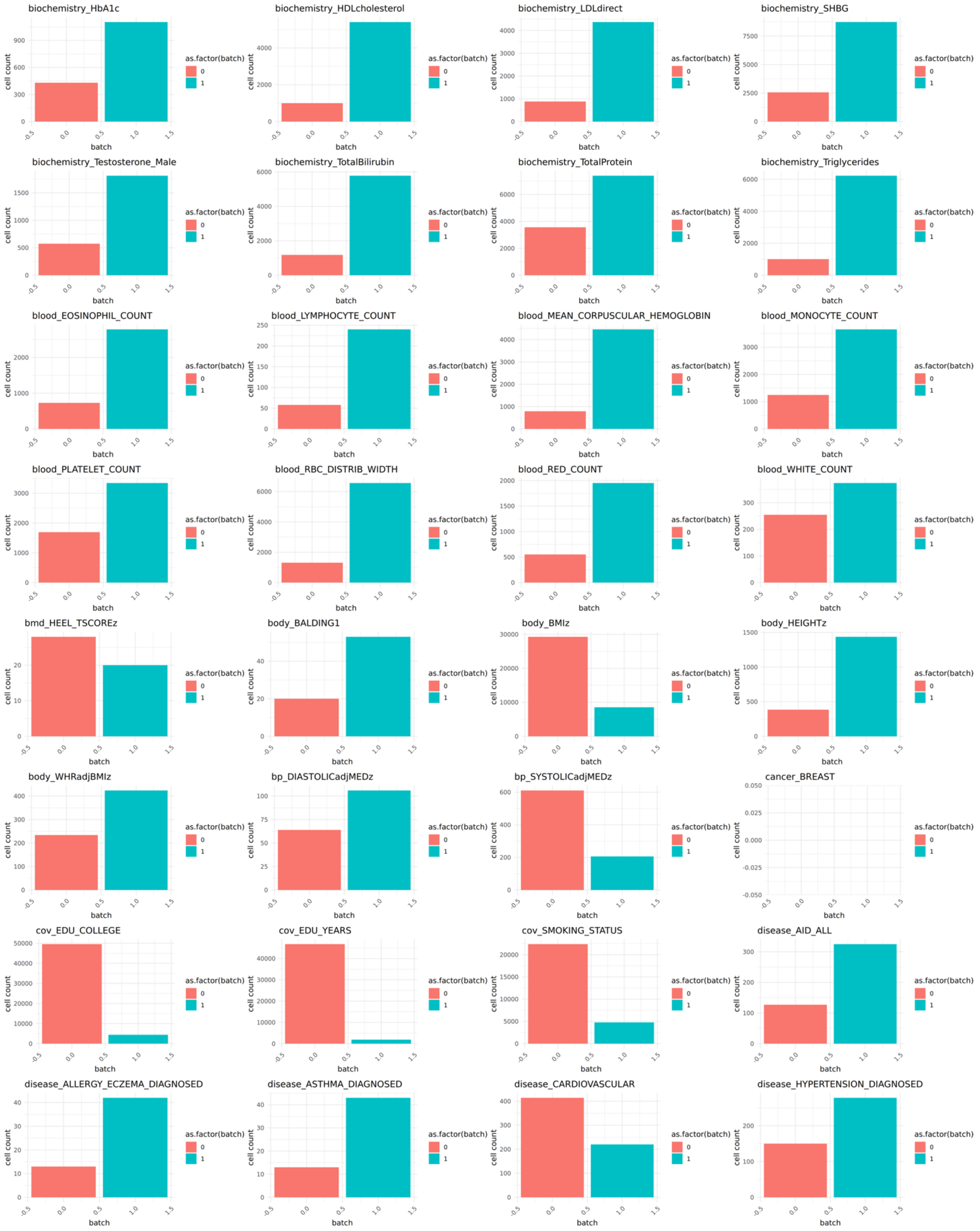


**Supplementary figure 9: Number of significant cells colored by batch (continue)**

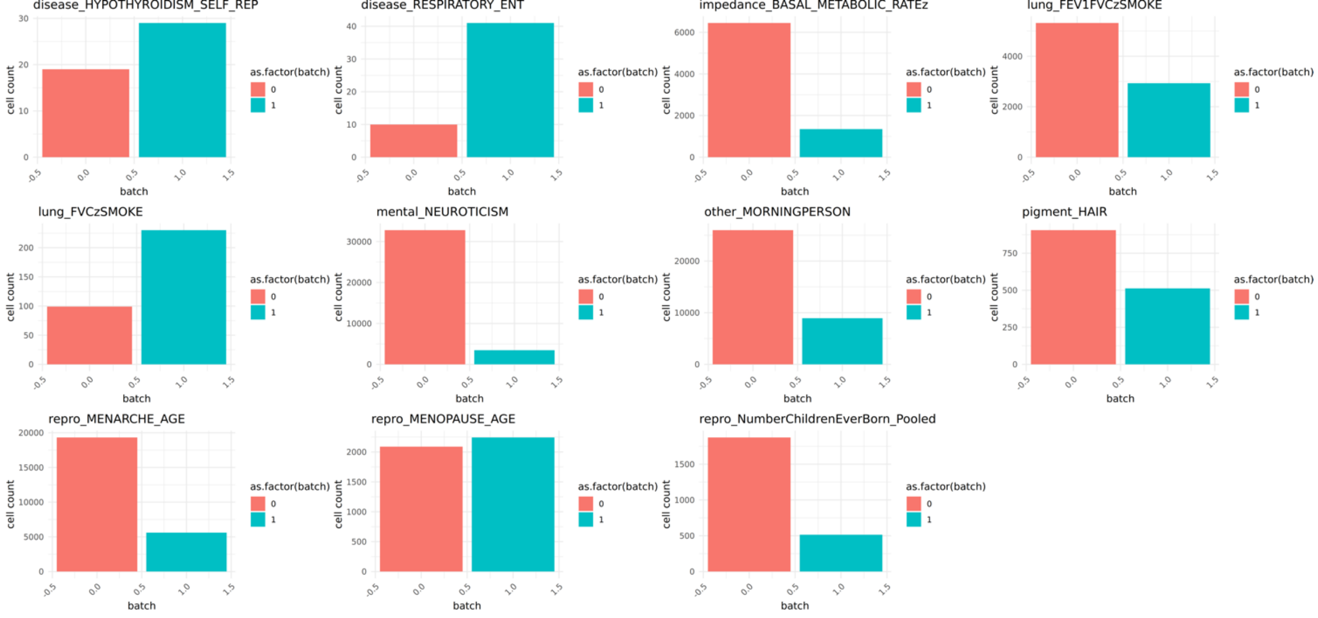


### Supplementary figure 9: Number of significant cells colored by batch (continue)

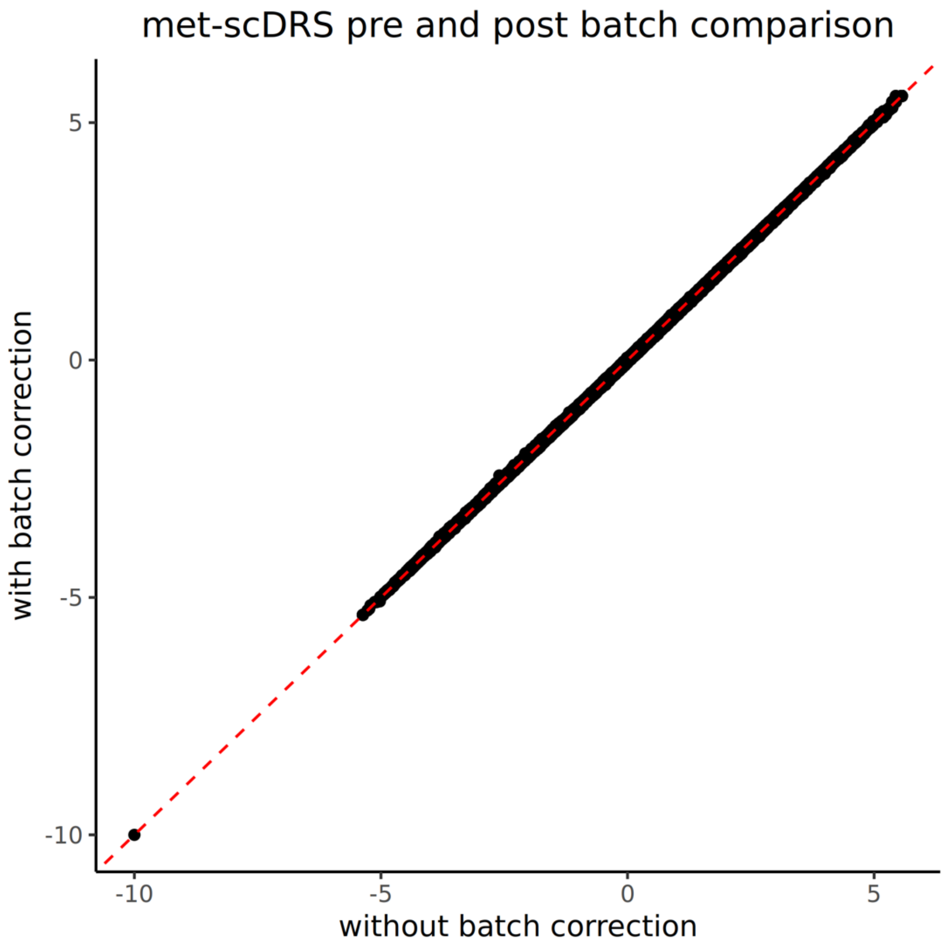


**Supplementary figure 10: Scatter plot visualizing met-scDRS before and after batch correction in a subset of GSE215353 data (n cells = 74777). The red dotted line represent X = Y line. Each dot represents a cell in the dataset.**

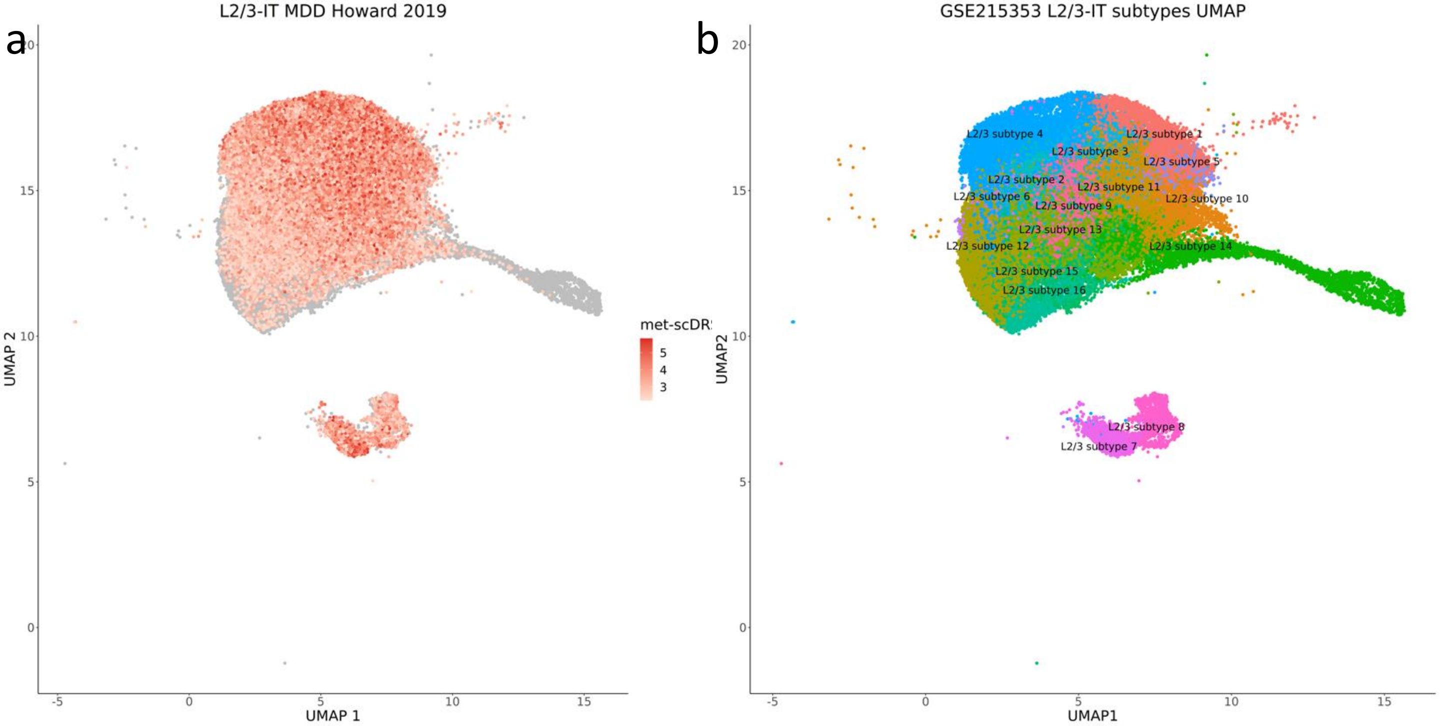


**Supplementary figure 11: a) UMAP plot colored by met-scDRS of L2/3–IT neurons for MDD and b) UMAP plot colored by subtype within excitatory L2/3 – IT neurons**

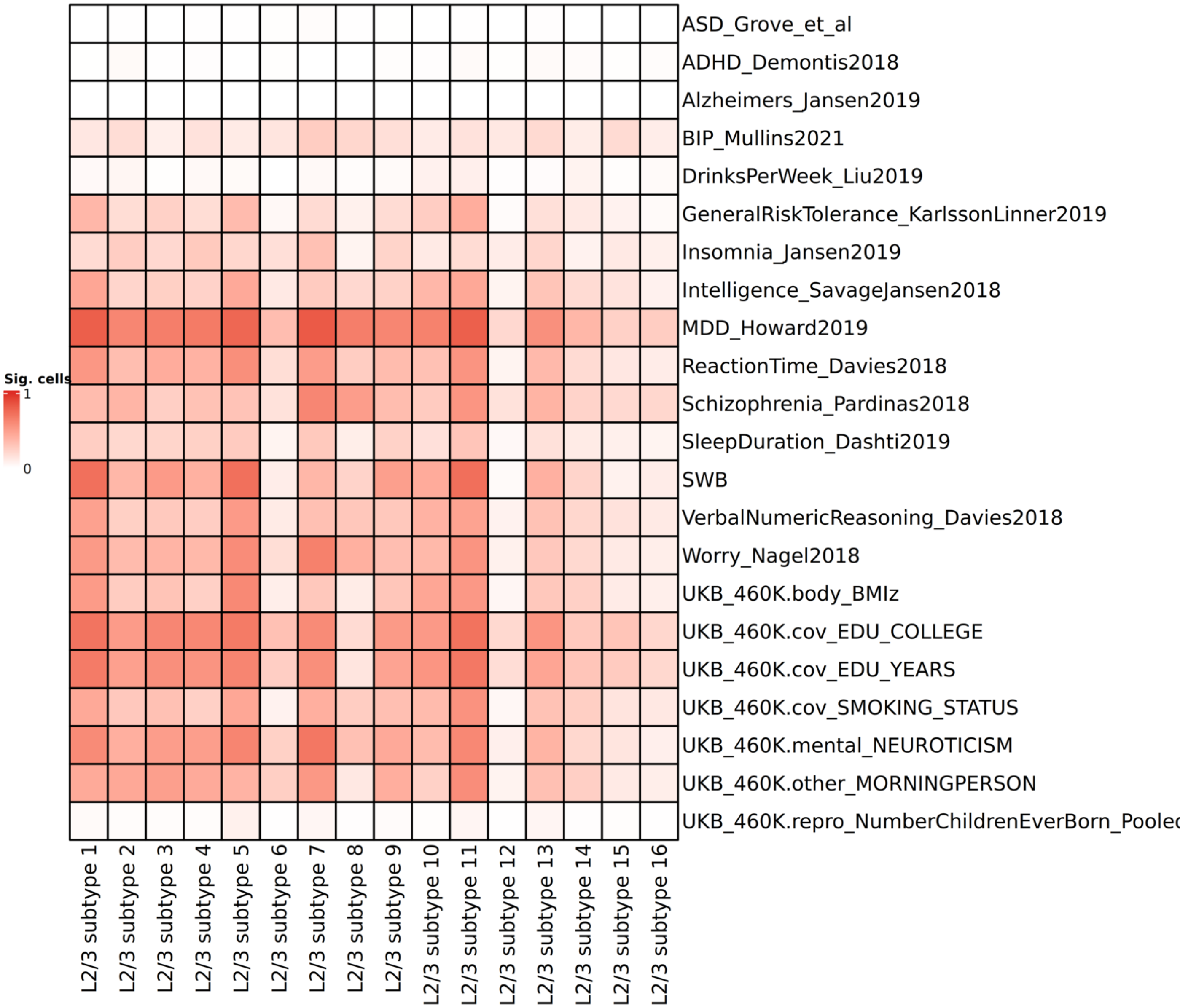


**Supplementary figure 12: Heatmap showing the proportion of cells in each of the L2/3-IT neuronal subtypes that are significant in each of the traits.**

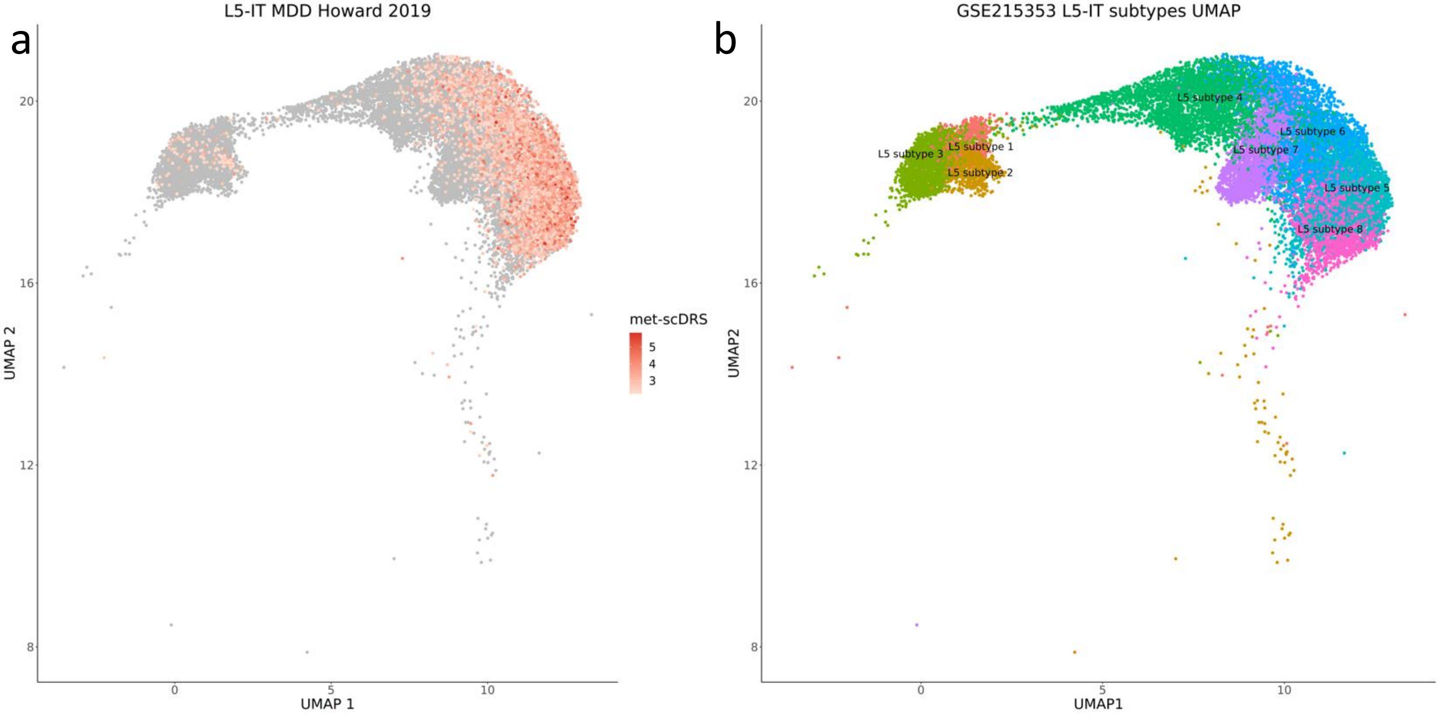


**Supplementary figure 13: a) UMAP plot colored by met-scDRS of L5–IT neurons for MDD and b) UMAP plot colored by subtype within excitatory L5 – IT neurons**

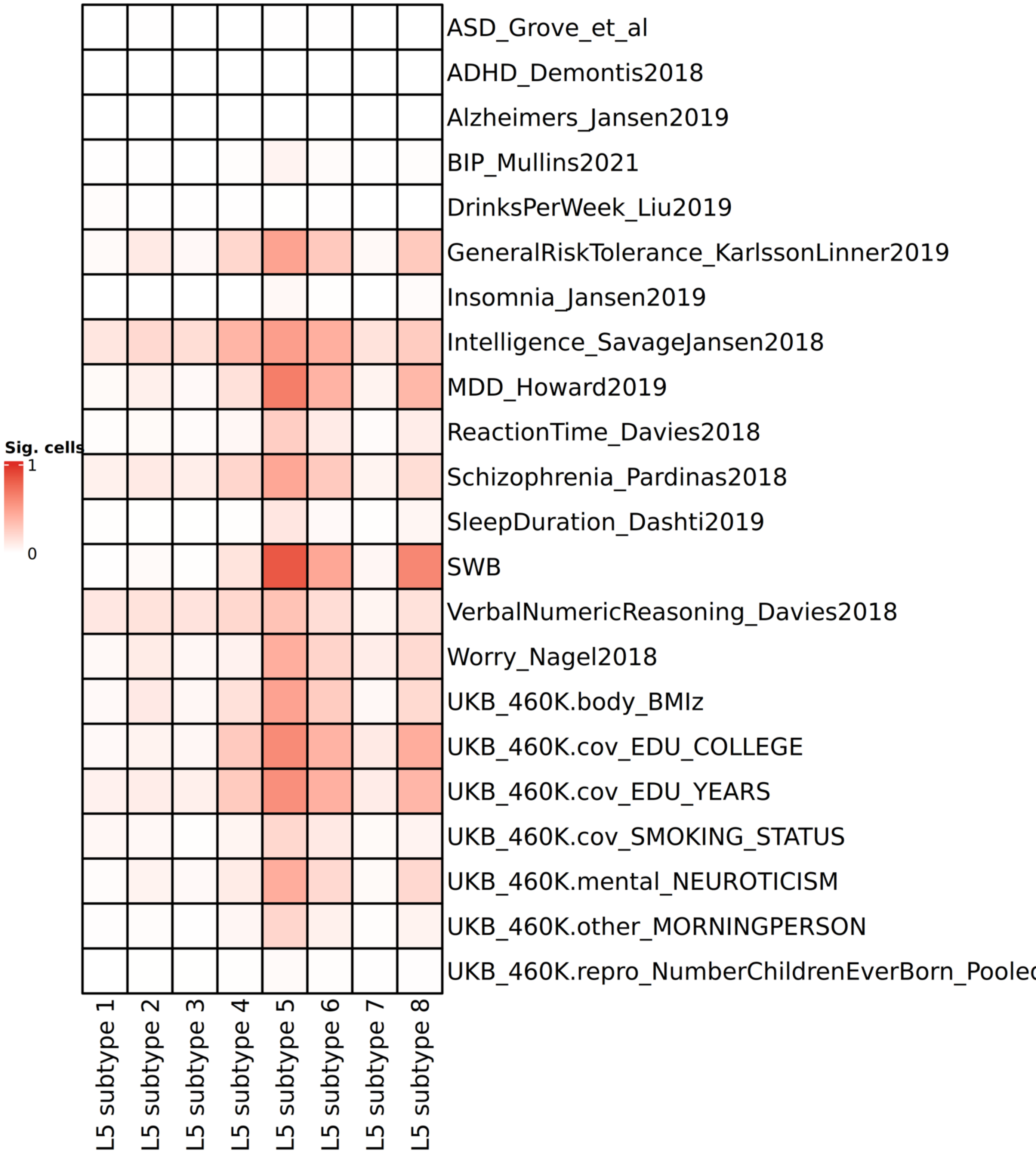


**Supplementary figure 14: Heatmap showing the proportion of cells in each of the L5-IT neuronal subtypes that are significant in each of the traits.**

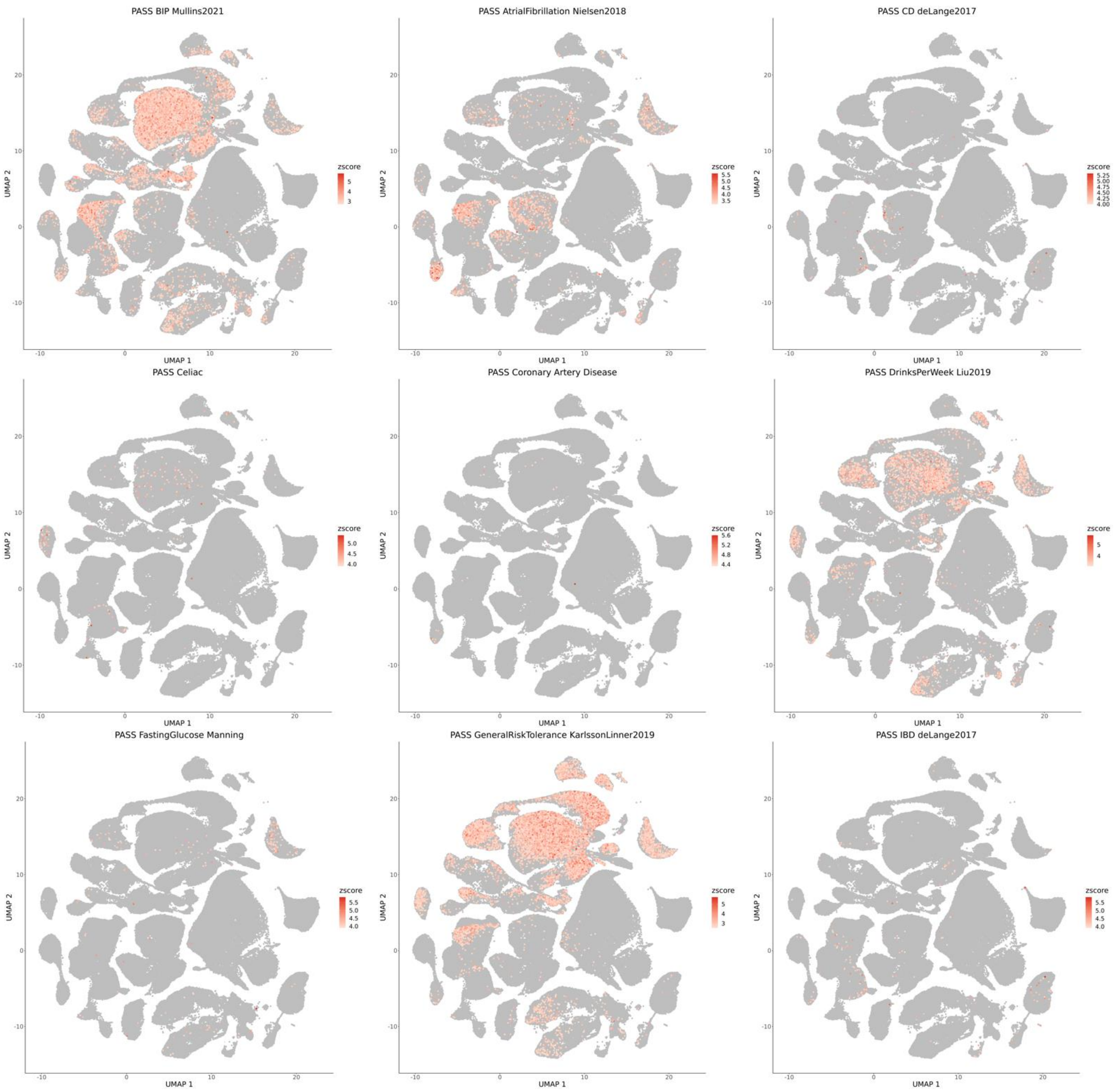


**Supplementary figure 15:** UMAP visualization using all cells in GSE215353 for all analyzed traits colored by non-CpG met-scDRS. The significant cells (FDR < 0.1) are colored with a gradient from white to red and the non-significant cells are colored as grey.

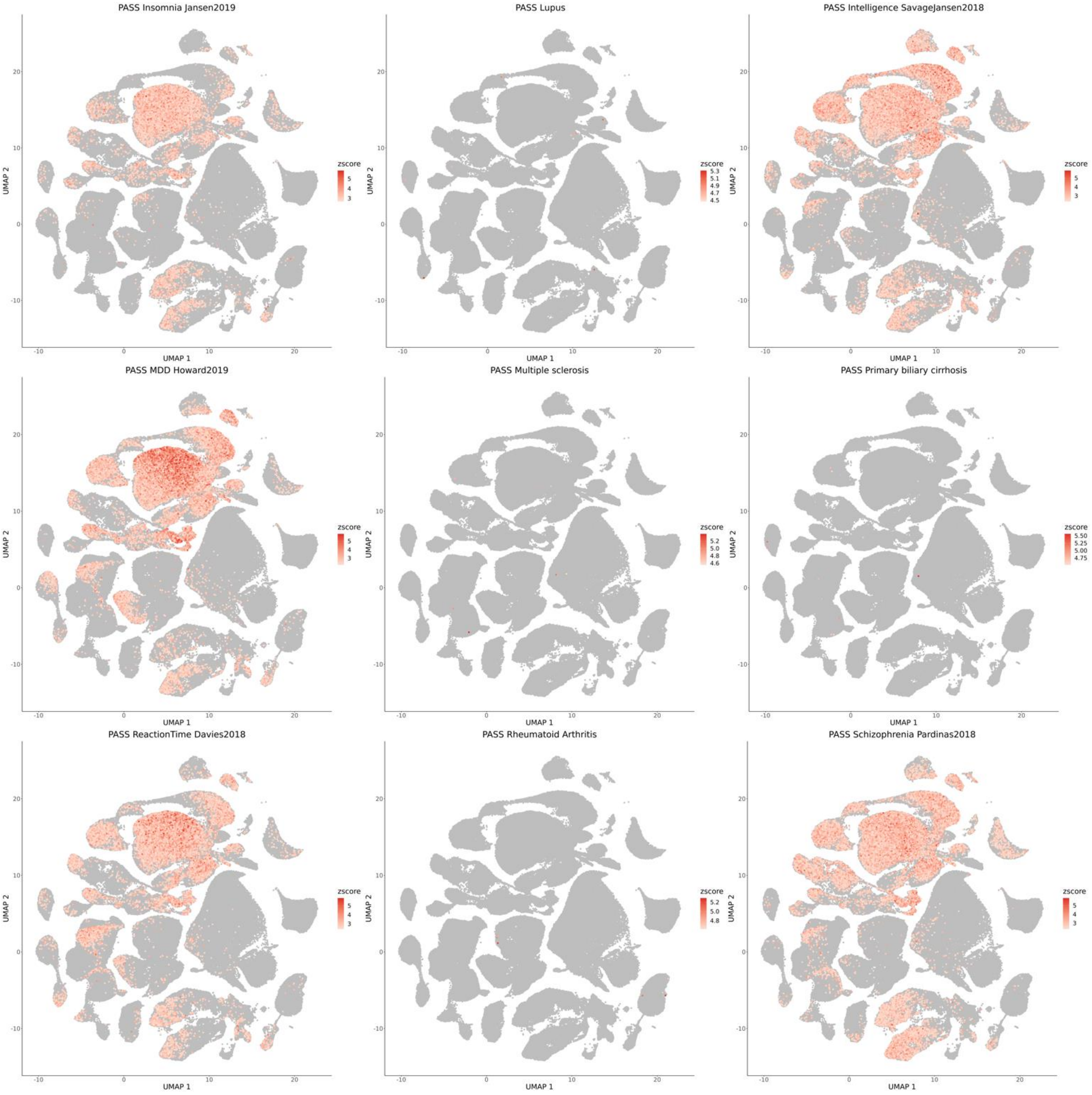


**Supplementary figure 15:** UMAP visualization using all cells in GSE215353 for all analyzed traits colored by non-CpG met-scDRS. The significant cells (FDR < 0.1) are colored with a gradient from white to red and the non-significant cells are colored as grey. (Continued)

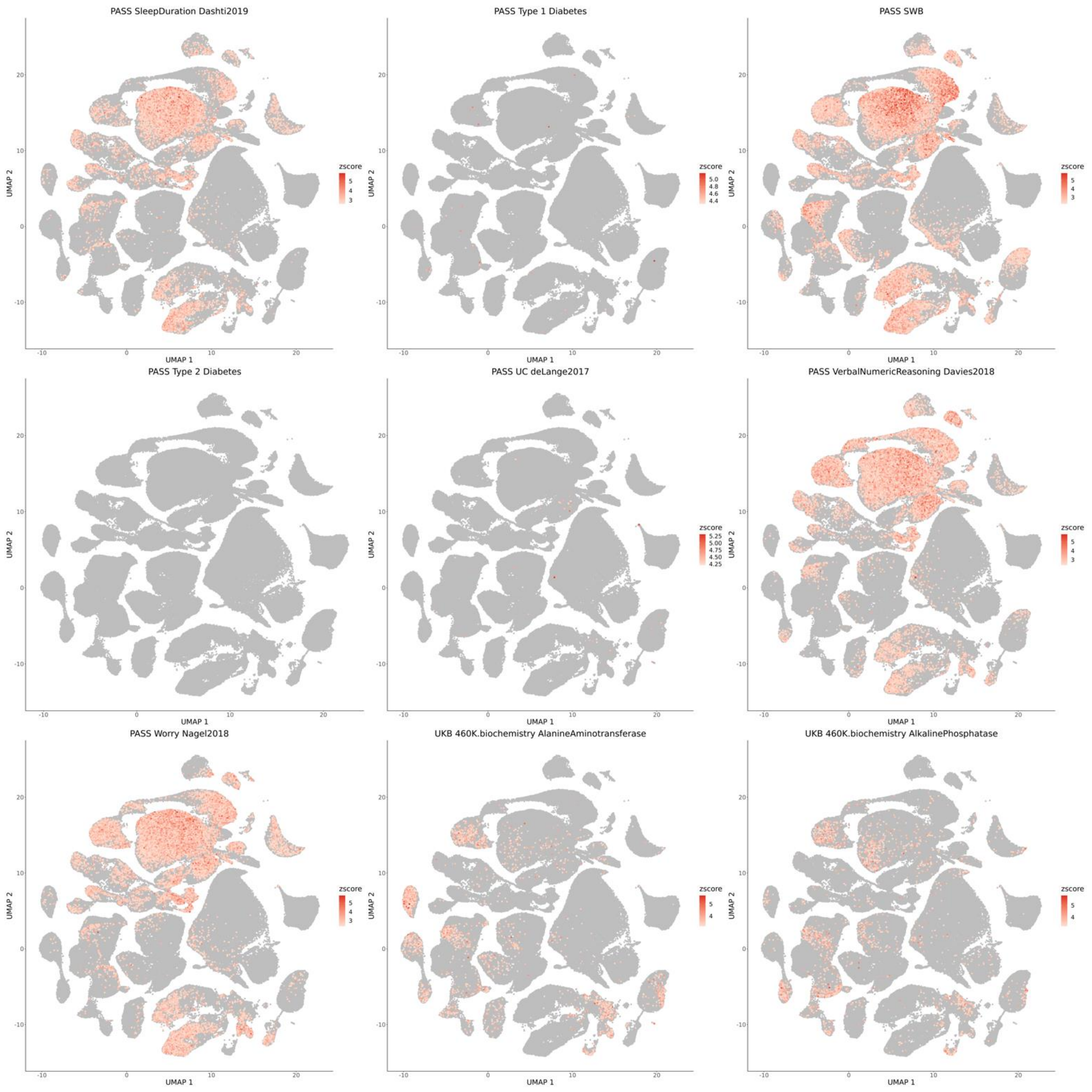


**Supplementary figure 15:** UMAP visualization using all cells in GSE215353 for all analyzed traits colored by non-CpG met-scDRS. The significant cells (FDR < 0.1) are colored with a gradient from white to red and the non-significant cells are colored as grey. (Continued)

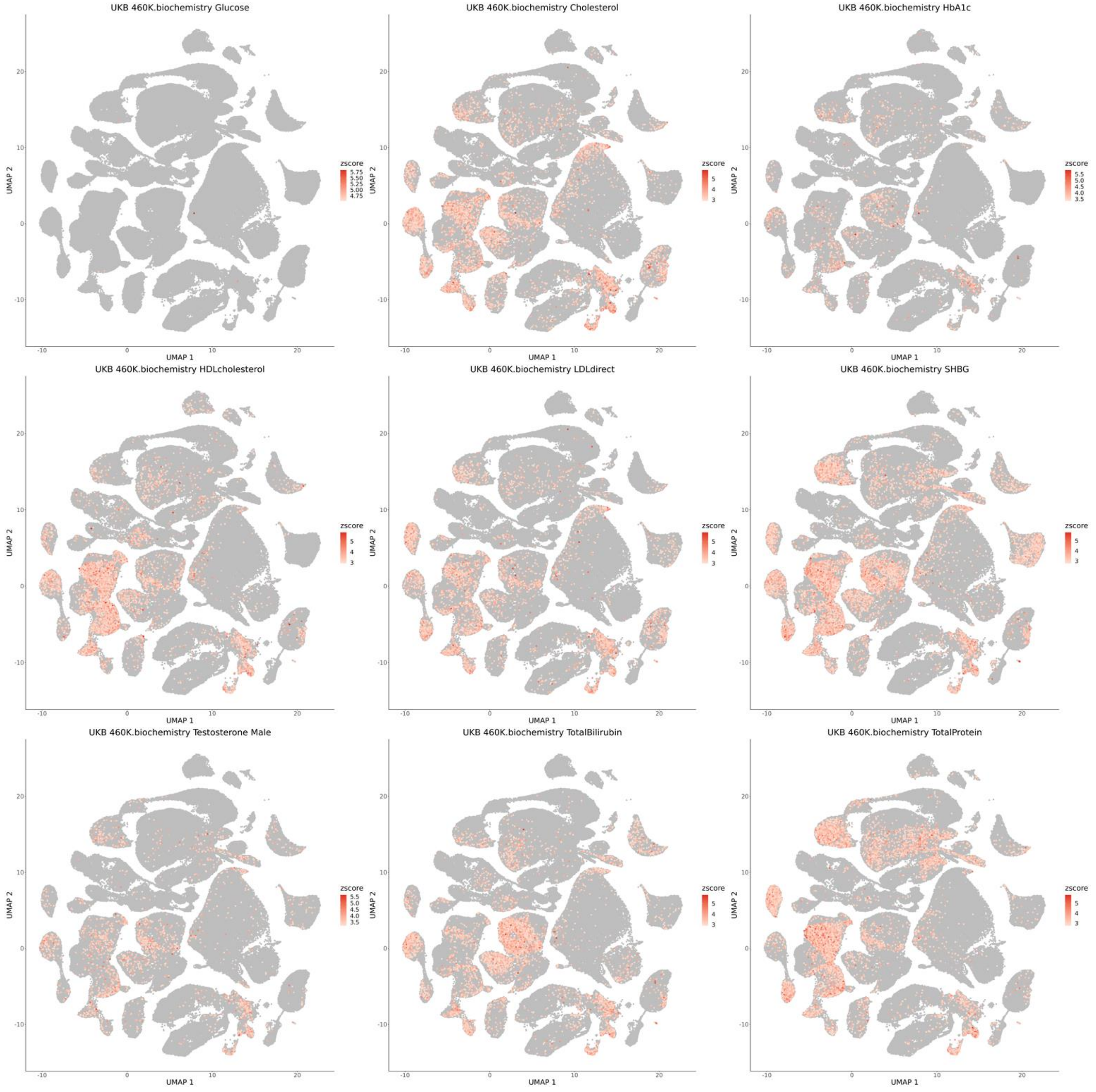


**Supplementary figure 15:** UMAP visualization using all cells in GSE215353 for all analyzed traits colored by non-CpG met-scDRS. The significant cells (FDR < 0.1) are colored with a gradient from white to red and the non-significant cells are colored as grey. (Continued)

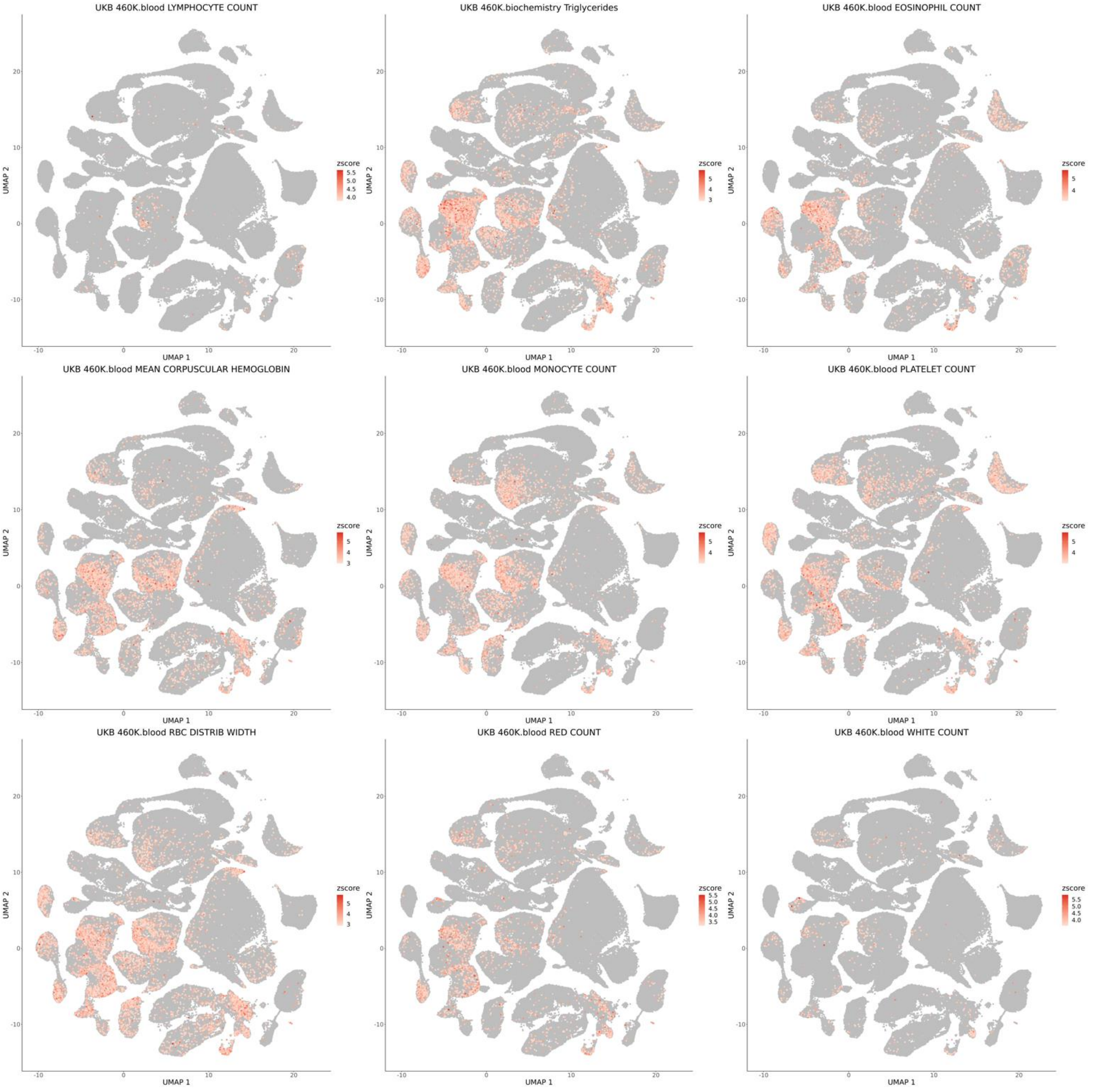


**Supplementary figure 15:** UMAP visualization using all cells in GSE215353 for all analyzed traits colored by non-CpG met-scDRS. The significant cells (FDR < 0.1) are colored with a gradient from white to red and the non-significant cells are colored as grey. (Continued)

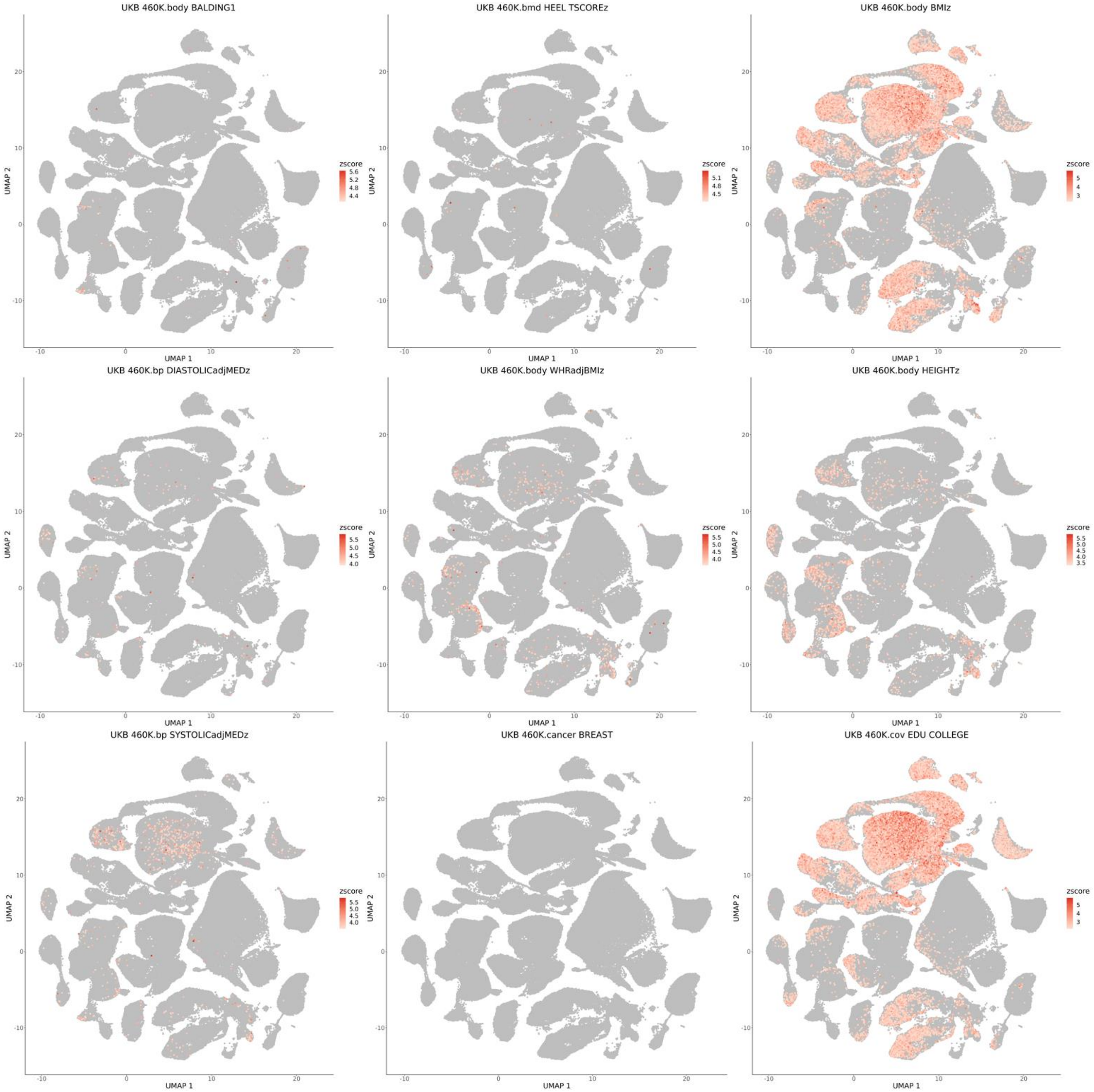


**Supplementary figure 15:** UMAP visualization using all cells in GSE215353 for all analyzed traits colored by non-CpG met-scDRS. The significant cells (FDR < 0.1) are colored with a gradient from white to red and the non-significant cells are colored as grey. (Continued)

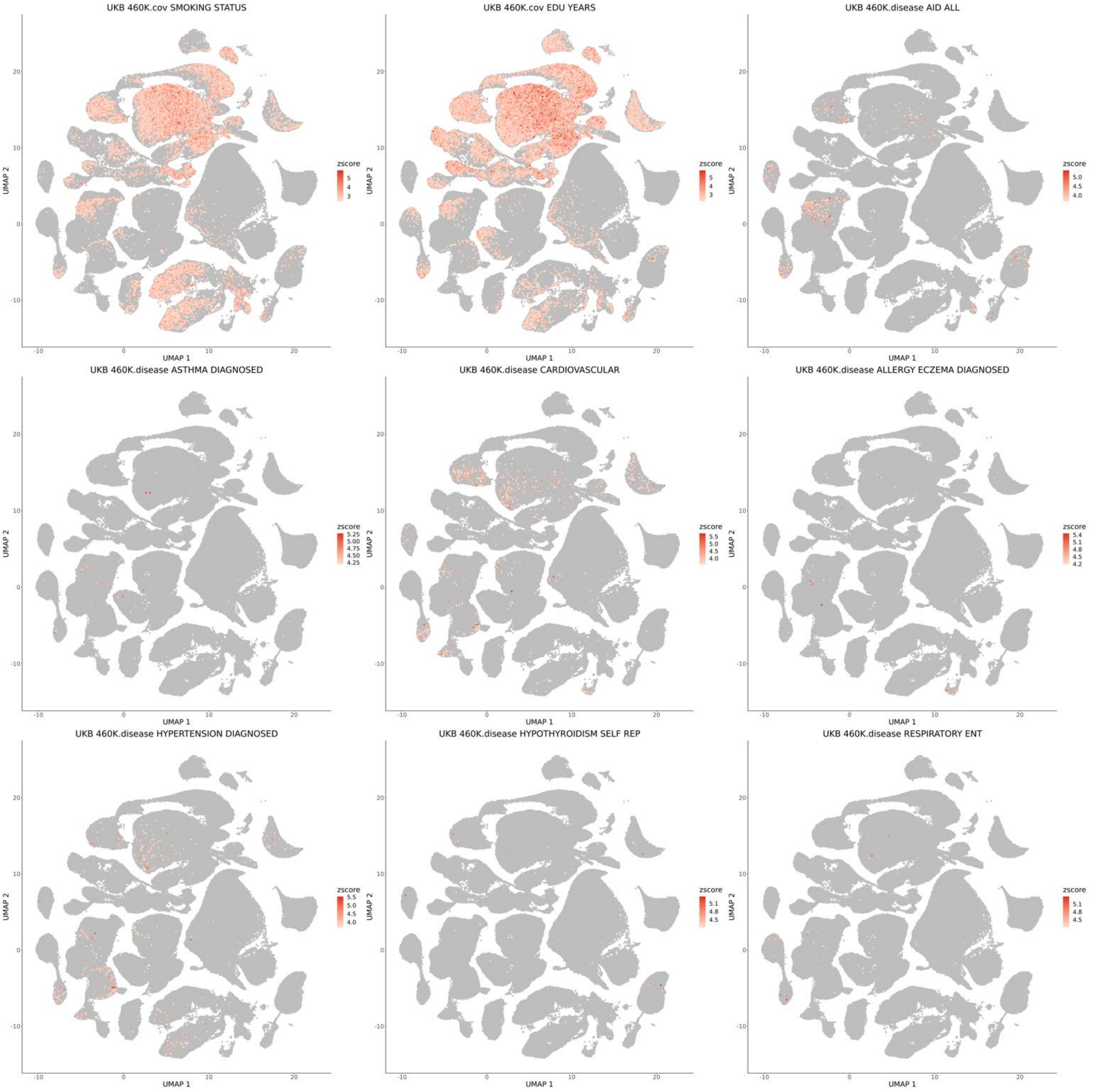


**Supplementary figure 15:** UMAP visualization using all cells in GSE215353 for all analyzed traits colored by non-CpG met-scDRS. The significant cells (FDR < 0.1) are colored with a gradient from white to red and the non-significant cells are colored as grey. (Continued)

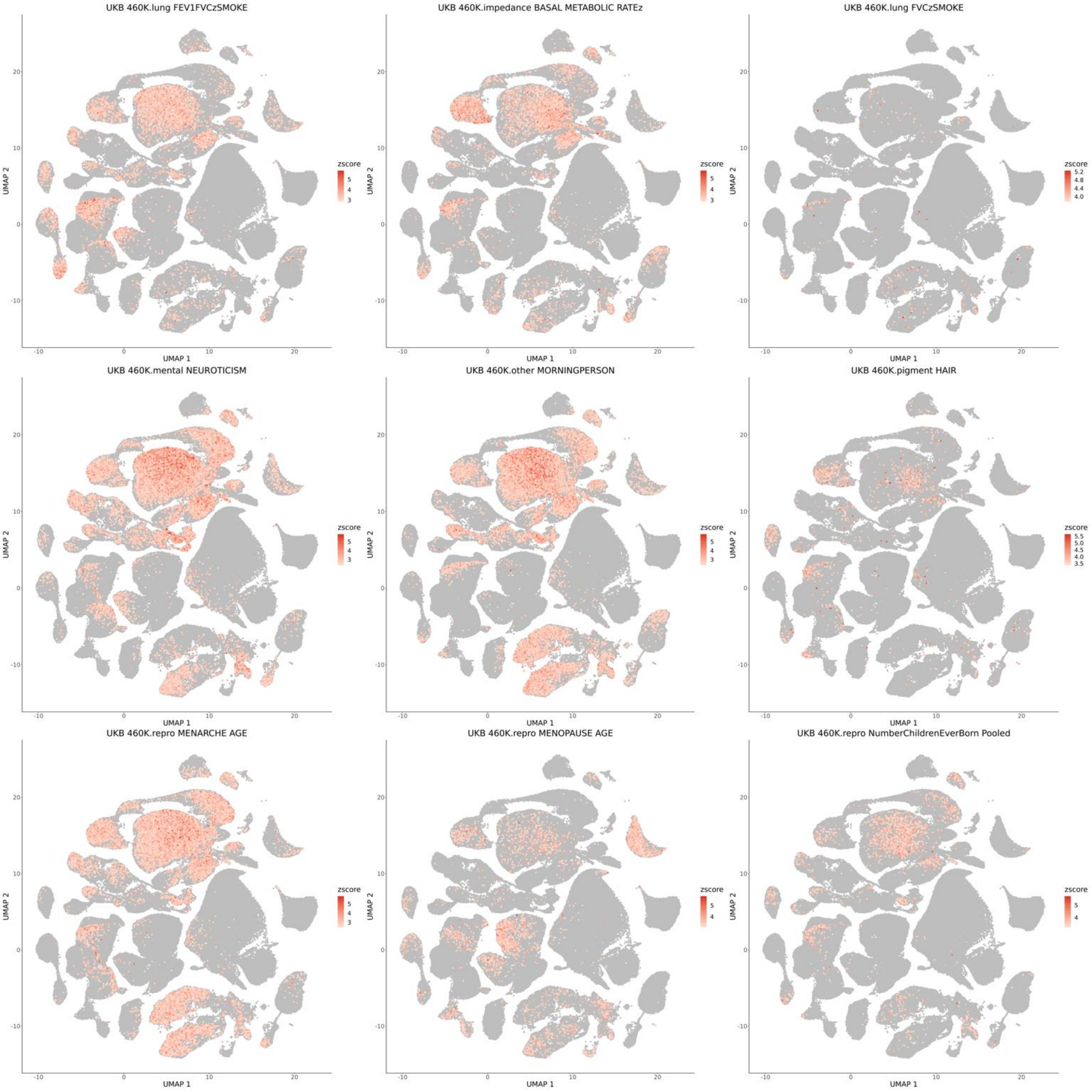


**Supplementary figure 15:** UMAP visualization using all cells in GSE215353 for all analyzed traits colored by non-CpG met-scDRS. The significant cells (FDR < 0.1) are colored with a gradient from white to red and the non-significant cells are colored as grey. (Continued)

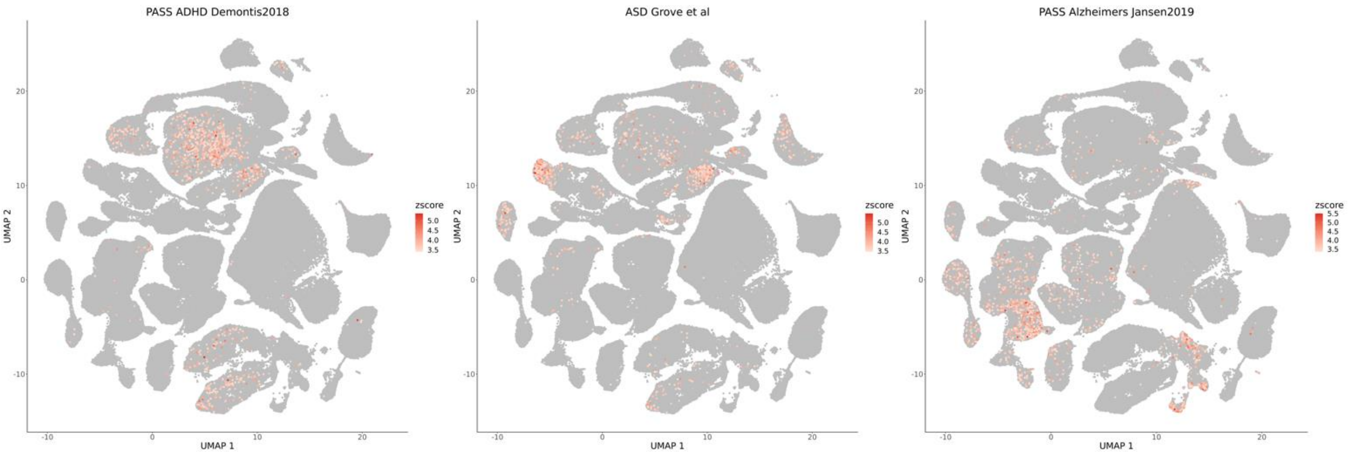


**Supplementary figure 15:** UMAP visualization using all cells in GSE215353 for all analyzed traits colored by non-CpG met-scDRS. The significant cells (FDR < 0.1) are colored with a gradient from white to red and the non-significant cells are colored as grey. (Continued)

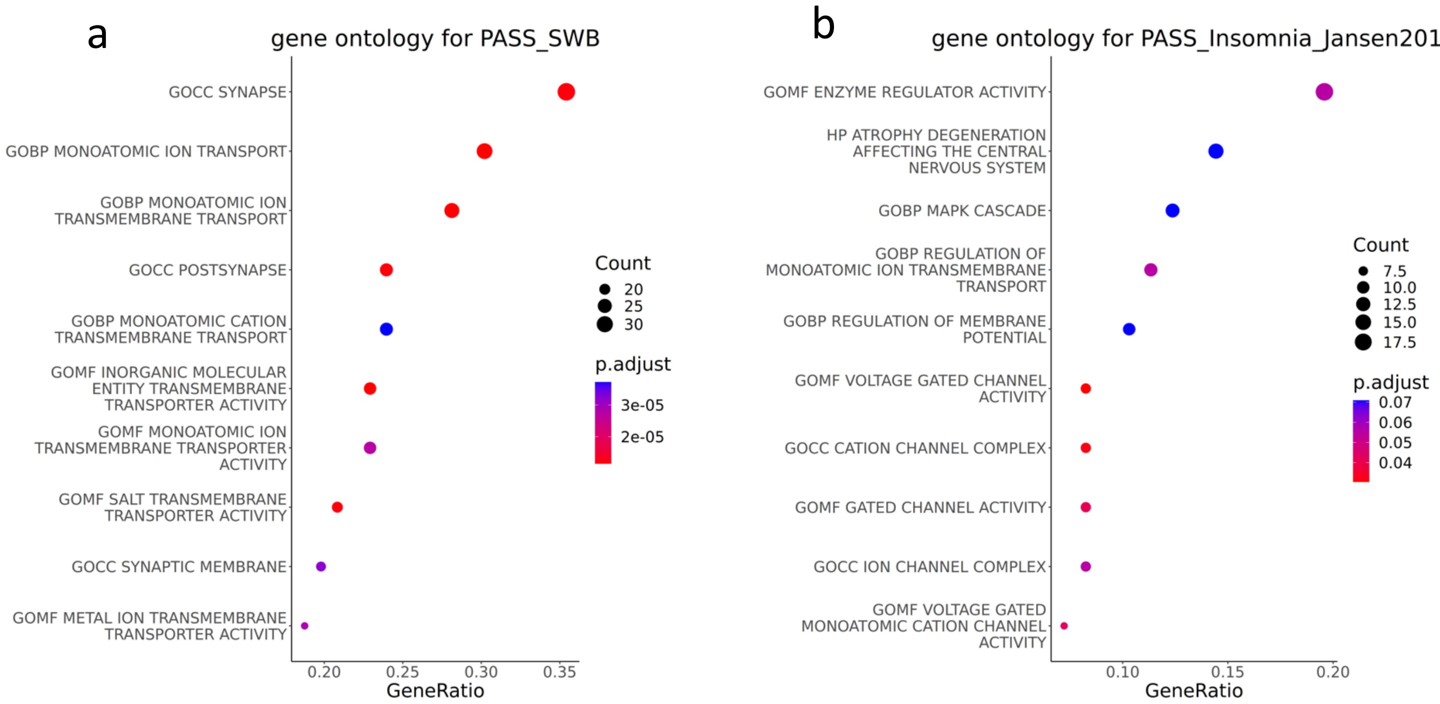


**Supplementary figure 16:** Gene ontology analysis using met-scDRS prioritized gene sets in a) social well being and b) Insomnia. For each of the pathways in Y axis, X axis shows the gene ratio between foreground and background; each dot is colored by FDR adjusted p value and sized by the number of genes in foreground that belong in that pathway

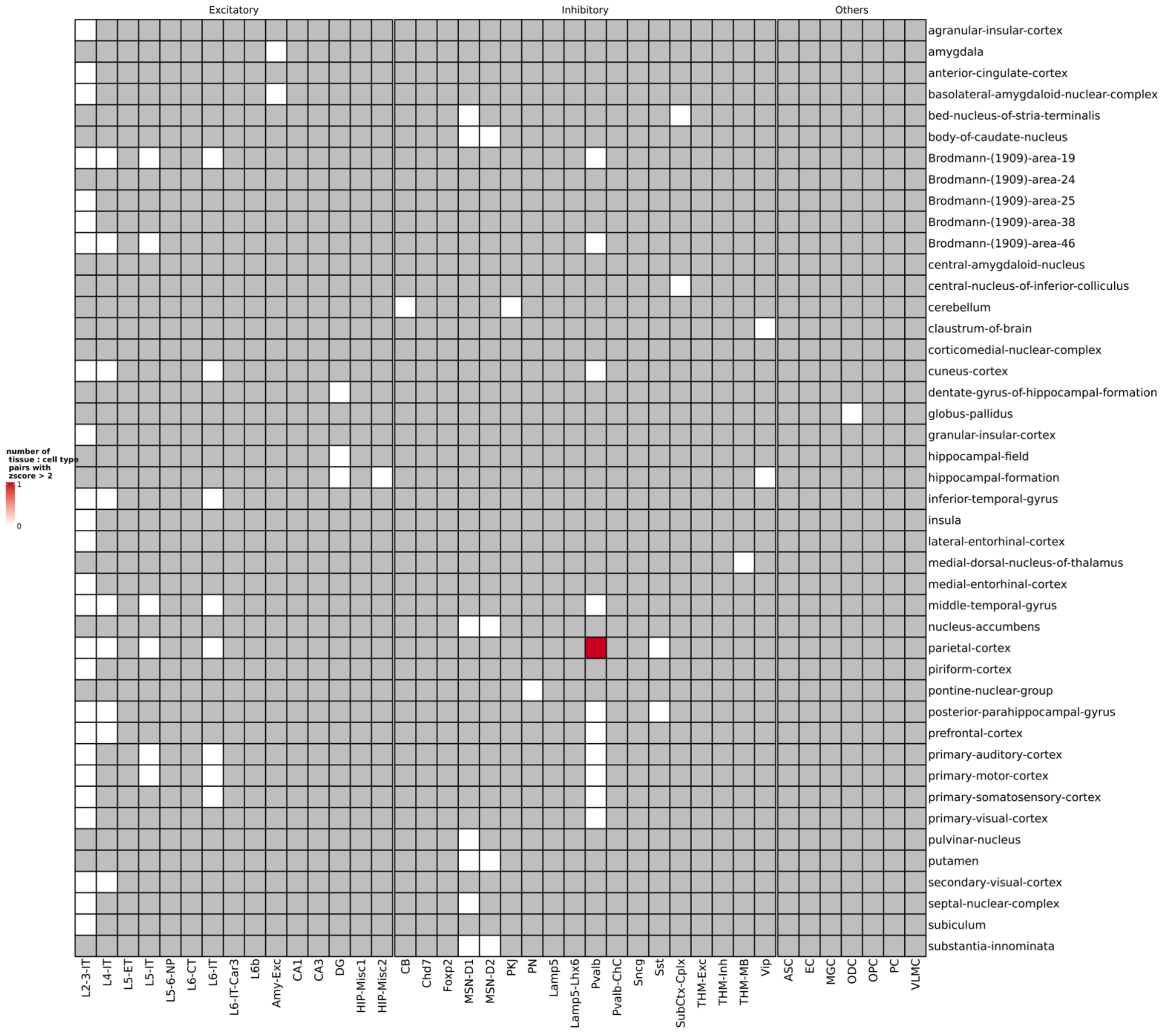


**Supplementary figure 17:** Heatmap visualization on number of non brain traits with > 2 specificity score in each of the tissue cell type pairs. Grey color represents not tested due to insufficient number of cells for testing in that tissue – cell type pair. White represents 0 traits that having specificity score > 2. and red color represent there is one trait with specificity score > 2.

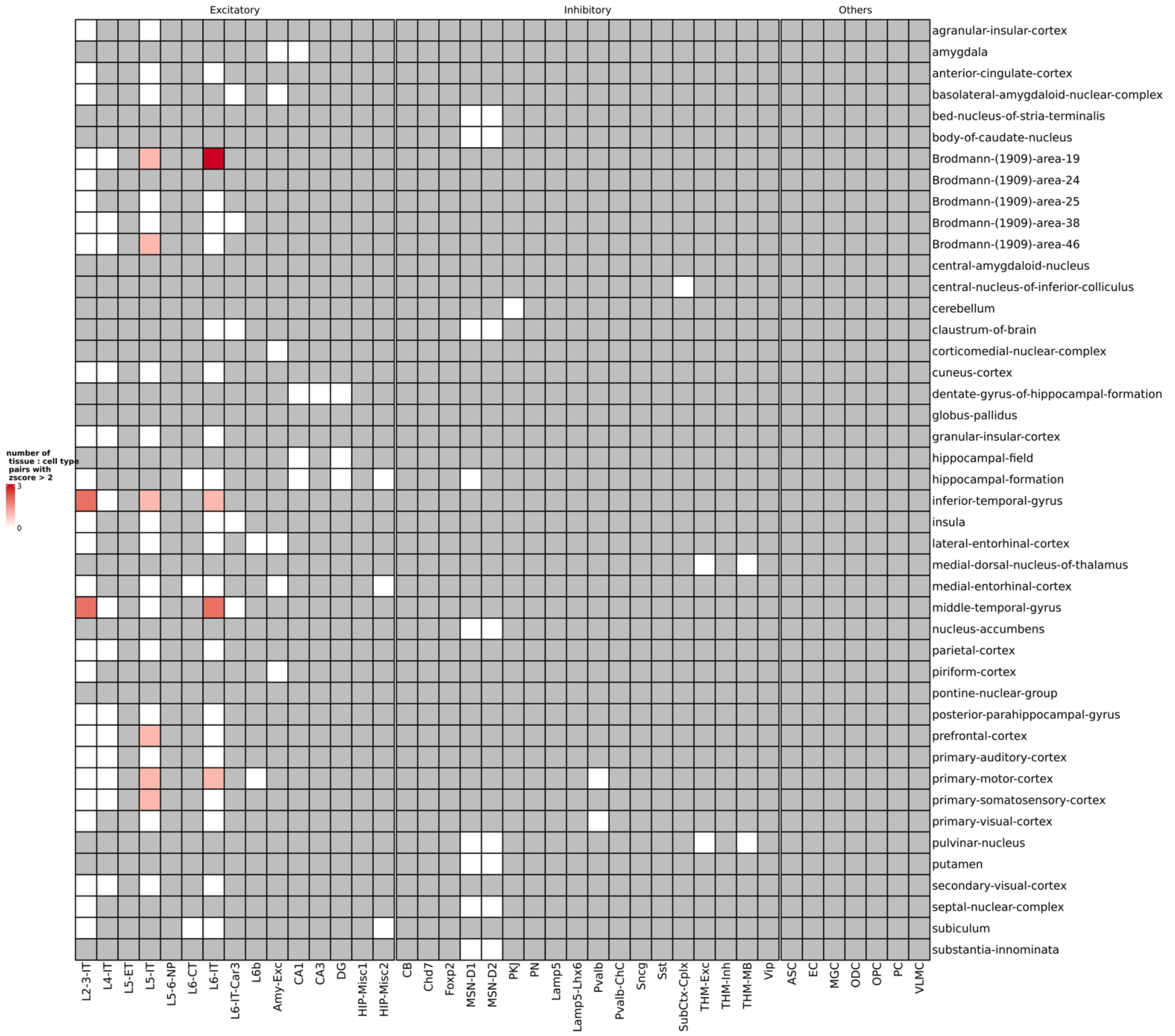


**Supplementary figure 18:** Heatmap visualization on number of brain traits with > 2 specificity score in each of the tissue cell type pairs. Grey color represents not tested due to insufficient number of cells for testing in that tissue – cell type pair. White represents 0 traits that having specificity score > 2. the number of traits is represented with a gradient from pink to red.

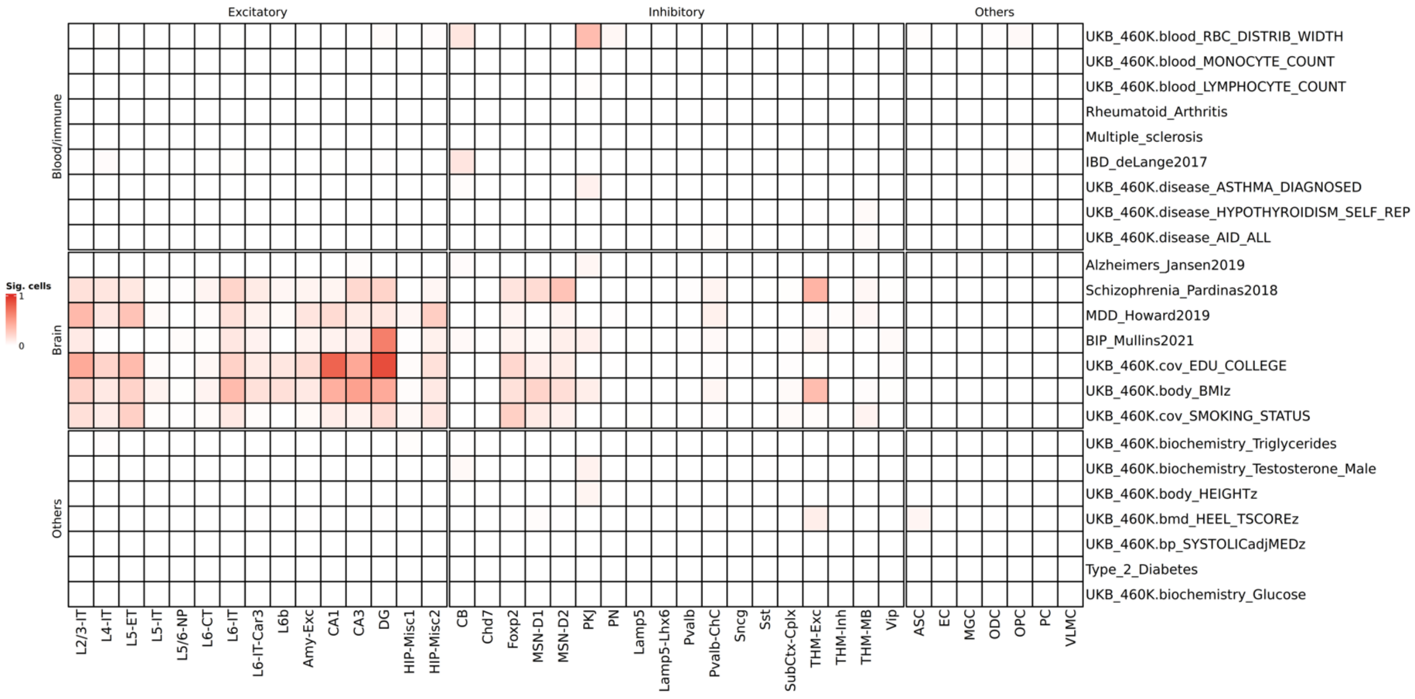


**Supplementary figure 19: Heatmap showing the proportion of cells that are significant to each traits in each of the neuronal major type computed using CpG methylation modality for GSE215353 subset data. i.e.: cells in cell type that are significant / number of cells in each cell type. The traits in the rows are selected to represent blood / immune, brain, and other traits.**

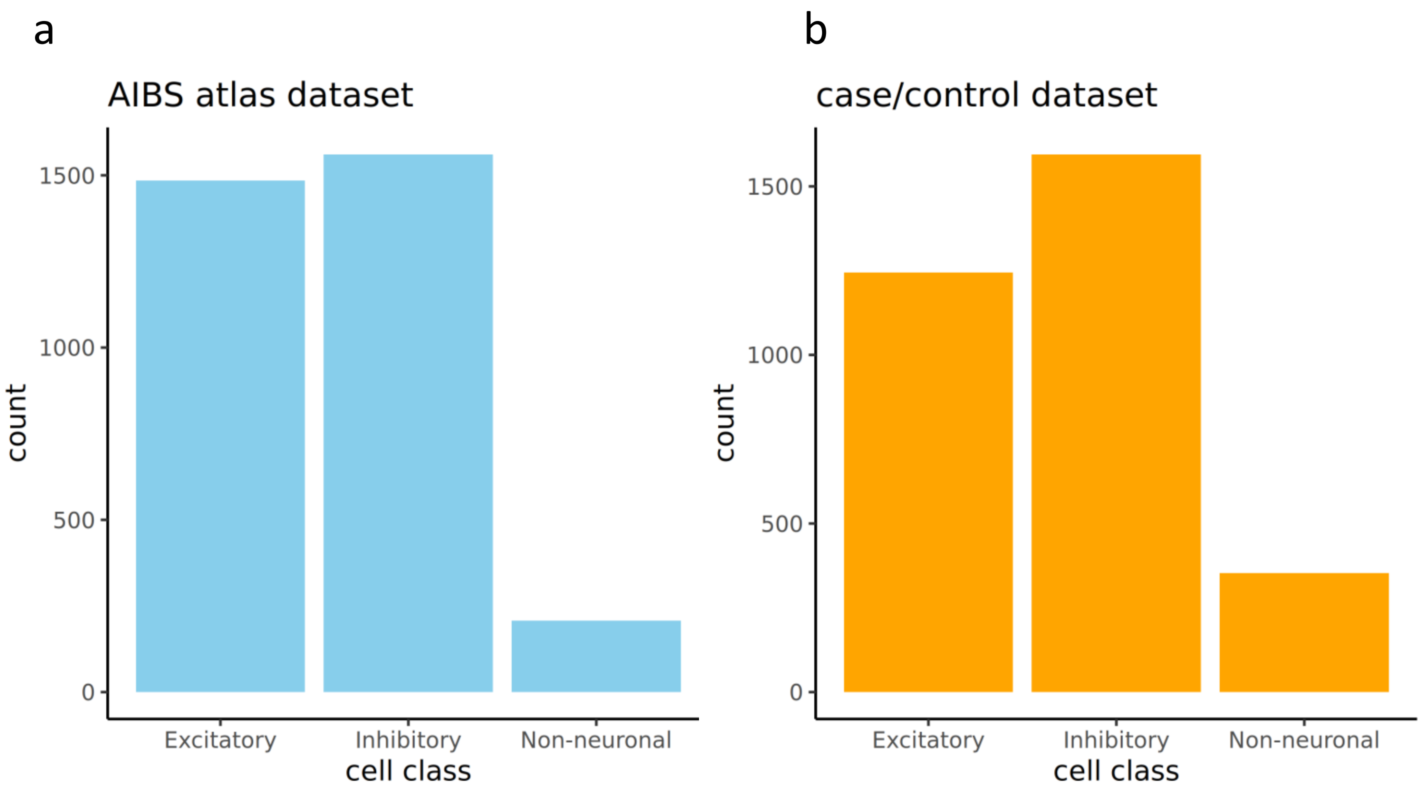


**Supplementary figure 20:** Bar chart indicating (a) number of marginally significant cells (uncorrected $p < 0.05$) that in the AIBS atlas dataset across cell class and (b) number of marginally significant cells in the case/ control dataset across cell class

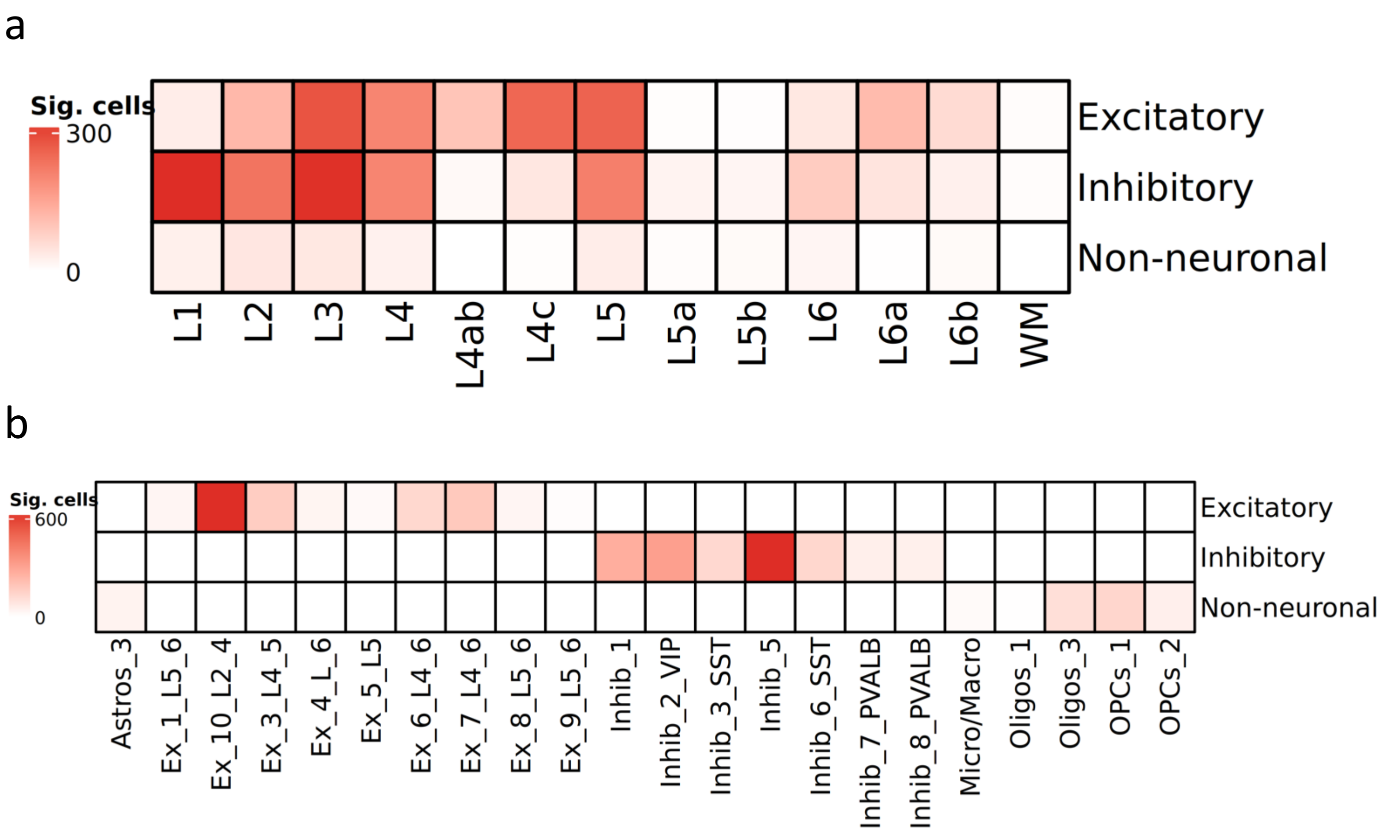


**Supplementary figure 21: (a)**Heatmap visualization on number of significant cells that are marginally significant (uncorrected $p < 0.05$) in AIBS transcriptomic dataset using MDD MAGMA genes with scDRS across layer and class. (**b**) Heatmap visualization on number of significant cells that are marginally significant in case/control dataset using MDD MAGMA genes with scDRS across cell type and class.
